# Glycosylation enzyme mRNA expression in dorsolateral prefrontal cortex of elderly patients with schizophrenia: Evidence for dysregulation of multiple glycosylation pathways

**DOI:** 10.1101/369314

**Authors:** Toni M. Mueller, Micah S. Simmons, Ashley T. Helix, Vahram Haroutunian, James H. Meador-Woodruff

## Abstract

Altered protein post-translational modifications such as glycosylation have become a target of investigation in the pathophysiology of schizophrenia. Disrupted glycosylation associated processes including atypical sphingolipid metabolism, reduced polysialylation of cell adhesion molecules, abnormal proteoglycan expression, and irregular glycan synthesis and branching have also been reported in this disorder. These pathways are regulated by the expression of glycosidases and glycosyltransferases, classes of enzymes which comprise approximately 2% of the genome. Many glycosylation enzymes can participate in multiple glycosylation pathways and dysregulation of enzyme expression could represent a common mechanism leading to a variety of glycan processing deficits in schizophrenia. In matched pairs of elderly schizophrenia and comparison subject (N = 12 pairs) dorsolateral prefrontal cortex, we measured mRNA levels of 84 key glycosylation enzymes by qPCR array. We found dysregulated transcript expression of 36 glycosylation enzymes from 12 functional categories. All of the abnormally expressed enzymes demonstrated increased transcript expression in schizophrenia, and many altered enzymes are known to modify substrates that have been previously implicated in the pathophysiology this illness. These findings suggest that abnormal glycosylation enzyme expression in schizophrenia may contribute to dysregulation of multiple glycosylation pathways, and disruptions of these central cell signaling processes may underlie a variety of deficits in schizophrenia.

---

The pathophysiology of schizophrenia is complex and remains incompletely understood; dysregulation across multiple brain regions, cell types, and neurotransmitter systems has been implicated in this disorder (Coyle, 1996; Davis et al., 1991; Harrison, 2000; McCullumsmith et al., 2003; Nakazawa et al., 2012). Neurotransmission abnormalities are well-documented in this illness, but the mechanisms underlying these irregularities have not been conclusively identified. Examination of gene and protein expression of neurotransmitter receptors and associated proteins have been often contradictory (Harrison and Weinberger, 2005; Kristiansen et al., 2007; Meador-Woodruff and Healy, 2000), while studies of posttranslational protein modifications (PTMs) have only recently become a target of investigation (Harrison, 2011, 2000; McCullumsmith and Meador-Woodruff, 2011). Alterations of central cell signaling processes mediated by PTMs may provide a common mechanism to explain disruptions involving multiple cell-types and neurotransmitter systems. Our lab has previously hypothesized that deficits of neurotransmission are not only due to abnormal expression of neurotransmitter-associated molecules, but may also result from abnormal receptor targeting and subcellular localization (Hammond et al., 2011; Mueller et al., 2015). Glycosylation is a PTM that plays a pivotal role in protein trafficking and subcellular targeting and has been shown in multiple studies to be abnormal in schizophrenia (Berretta, 2012; Ikemoto, 2014; Mueller et al., 2014; Narayan et al., 2009; Pantazopoulos et al., 2013; Sato and Kitajima, 2013; Stanta et al., 2010; Tucholski et al., 2013b, 2013a).

Glycosylation is an enzyme-mediated process whereby a carbohydrate is attached to a protein, lipid, or glycan substrate. There are multiple glycosylation pathways in the cell, including N-linked and O-linked protein glycosylation, sphingolipid metabolism, and mannose-6-phosphate catabolism. We have previously reported abnormal N-linked glycosylation in schizophrenia of glutamate transporters as well as subunits from α-amino-3-hydroxy-5-methyl-4-isoxazolepropionic acid (AMPA), kainate, and GABAA receptor families (Bauer et al., 2010; Mueller et al., 2014; Tucholski et al., 2013b, 2013a). We have demonstrated that receptors containing abnormally N-glycosylated subunits also exhibit abnormal subcellular distribution in schizophrenia (Hammond et al., 2010; Mueller et al., 2015), suggesting cellular consequences of abnormal protein glycosylation. In addition to the N-linked glycosylation alterations we have reported, abnormalities of glycan structures expressed in serum and cerebrospinal fluid (CSF) (Stanta et al., 2010), abnormal expression of genes associated with sphingolipid metabolism (Narayan et al., 2009), decreased polysialylation of cell adhesion molecules (Barbeau et al., 1995; Isomura et al., 2011), and abnormalities of chondroitin sulfate proteoglycan expression (Berretta, 2012; Pantazopoulos et al., 2013) have all been reported in schizophrenia. Together, these reports provide evidence that multiple types of glycan processing including synthesis and correct branching of glycan structures, post-translational glycosylation of protein substrates, synthesis and metabolism of glycolipid conjugates, and degradation of complex glycans may all be disrupted in schizophrenia. Although some enzymes are involved in only one type of glycosylation, others have diverse functions across multiple glycosylation pathways (DeMarco and Woods, 2008; Keith et al., 2005; Moremen et al., 2012; Ohtsubo and Marth, 2006; Röttger et al., 1998; Spiro, 2002).

Given reports of several types of glycosylation abnormalities and enzyme sharing between glycosylation pathways, it is likely that disruptions of glycan processing in schizophrenia are not exclusive to a particular glycosylation pathway but may reflect more widespread dysregulation. Based on our prior reports of abnormal N-glycosylation of neurotransmitter receptor subunits, we predicted that enzymes specifically associated with glycosylation in the endoplasmic reticulum (ER) would be abnormally expressed in schizophrenia. To investigate the hypothesis that multiple glycosylation pathways are disrupted due to abnormal expression of ER-localized enzymes in schizophrenia, we measured the transcript expression of 84 key glycosylation enzymes in the dorsolateral prefrontal cortex (DLPFC) of paired schizophrenia and comparison subjects.

## Materials and Methods

### Subjects and Tissue Preparation

Subjects were recruited by the Mount Sinai Medical Center Brain Bank. Assessment, consent, and postmortem procedures were conducted as required by the Institutional Review Boards of Pilgrim Psychiatric Center, Mount Sinai School of Medicine, and the Bronx Veterans Administration Medical Center. Subject medical history was assessed for history of psychiatric illnesses and history of drug or alcohol abuse. Patients had a documented history of psychiatric symptoms before age 40 and 10 or more years of hospitalization with a diagnosis of schizophrenia using DSM-III-R criteria as determined by two clinicians; comparison subjects had no documented history of psychiatric illnesses. Criteria for subject exclusion included a history of substance abuse, death by suicide, or coma for greater than 6 hours prior to death. Neither schizophrenia nor comparison subjects had evidence of neuropathological lesions or neurodegenerative disorders including Alzheimer’s disease (Powchik et al., 1993; Purohit et al., 1998). Whole brains were dissected into 1cm thick slabs. The full thickness of gray matter from the DLPFC (Brodmann 9/46) was cut into 1cm^3^ blocks from 12 pairs of schizophrenia and matched comparison subjects (Tables 1 and 2) and stored at - 80°C until sectioning. Tissue was thawed to -20°C to be sectioned by a microtome (AO Scientific Instruments) at 15 μm. Tissue sections were placed onto Fisher-brand Superfrost/Plus positively charged microscope slides (Fisher Scientific), air dried, and stored at -80°C.

**Table 1.** Summary of subject demographics.

|  | Comparison | Schizophrenia |
| --- | --- | --- |
| **n** | 12 | 12 |
| **Age** | 75.6 ± 9.4 | 75.8 ± 10.8 |
| **Sex** | 6F/6M | 6F/6M |
| **PMI (min)** | 600 ± 453 | 897 ± 395 |
| **Tissue pH** | 6.56 ± 0.29 | 6.45 ± 0.20 |
| **On/Off Rx** | 0/12 | 7/5 |

Abbreviations: Male (M), Female (F), Postmortem interval (PMI), Antipsychotic medication (Rx). Values are expressed as means ± standard deviation. Off Rx indicates patients that had not received antipsychotic medications for 6 weeks or more prior to death.

**Table 2.** Paired subjects.

|  |  | Comparison |  |  | Schizophrenia |  |  |  |
| --- | --- | --- | --- | --- | --- | --- | --- | --- |
|  |  |  |  | PMI |  |  | PMI |  |
| Pair | Sex | Age | pH | (hours) | Age | pH | (hours) | Rx |
| 1 | F | 73 | 6.98 | 3.0 | 70 | 6.51 | 13.2 | Off |
| 2 | F | 78 | 6.97 | 16.0 | 75 | 6.49 | 21.5 | Off |
| 3 | F | 81 | 6.37 | 19.4 | 81 | 6.67 | 15.1 | Off |
| 4 | F | 79 | 6.38 | 10.1 | 77 | 6.01 | 9.7 | On |
| 5 | F | 89 | 6.72 | 2.3 | 89 | 6.20 | 9.6 | On |
| 6 | F | 66 | 6.85 | 22.6 | 62 | 6.74 | 23.7 | On |
| 7 | M | 76 | 6.32 | 2.9 | 78 | 6.64 | 26.1 | Off |
| 8 | M | 75 | 6.43 | 5.0 | 80 | 6.37 | 15.4 | Off |
| 9 | M | 70 | 6.10 | 6.7 | 73 | 6.50 | 7.9 | On |
| 10 | M | 93 | 6.28 | 4.2 | 97 | 6.50 | 9.3 | On |
| 11 | M | 69 | 6.67 | 7.4 | 70 | 6.35 | 7.2 | On |
| 12 | M | 59 | 6.67 | 20.4 | 57 | 6.40 | 20.7 | On |

The comparison subject in pair 9 failed to amplify for > 20% of genes measured and was excluded from primary statistical analyses. Abbreviations: Male (M), Female (F), Postmortem interval (PMI), Antipsychotic medication (Rx). Off Rx indicates patients that had not received antipsychotic medications for 6 weeks or more prior to death.

### RNA Extraction and cDNA Pre-amplification

The full thickness of grey matter from DLPFC was scraped from 2 glass slides per subject using sterile razor blades and homogenized with a pestle in Picopure Extraction Buffer. RNA from each sample was isolated using a Picopure RNA Isolation kit (cat# KIT0214, Life Technologies). RNA concentration was determined by spectrophotometry at 260nm, and equal amounts of RNA for each subject were incubated with 1 unit of DNase I (Promega) per gram of RNA at 37 °C for 30 minutes then at 65 °C for 15 minutes. RNA was reverse transcribed to cDNA using a High Capacity cDNA Reverse Transcription Kit (cat# 4368813, Applied Biosystems), then pre-amplified using a RT^2^ Pre-AMP cDNA Synthesis Kit (cat# 330451, Qiagen) using the protocol provided and including the optional side reaction reducer.

### Comparative qRT-PCR

Pre-amplified cDNA for each subject was plated in a RT^2^ Human Glycosylation PCR Array 96-well plate (cat# PAHS-046A, Qiagen) and run on a Stratagene MX3000P QPCR System (Agilent Technologies, La Jolla, CA) at 50°C for 2 minutes, 95°C for 10 minutes, followed by 50 cycles of 95°C for 15 seconds and 60°C for 1 minute. Cycle threshold (Ct) values were obtained using Mx3000P v4.10 software (Stratagene, Agilent Technologies) with FAM threshold fluorescence = 1.000. Any wells that failed to amplify to the predefined threshold in 50 cycles (indicated as “No Ct” in instrument output) were excluded from primary statistical analysis. One comparison subject failed to amplify within 50 RT-PCR cycles in over 20% of the target genes and was excluded from statistical analysis.

### Statistical Analysis

Primary statistical analysis was performed on gene expression values. Sample size was determined *a priori* using a statistical power calculation (β = 0.2, π = 0.8) based on previous studies of gene expression from the lab. The investigators who executed the experiment were blind to subject diagnosis and subject pairing; the investigator who performed statistical analyses did not participate in the generation of data. B2M, HPRT1, RPL13A, GAPDH, and ACTB were used as housekeeping genes after verifying their expression was unchanged between diagnostic groups (Figure 1). The ∆Ct value for each subject was calculated by subtracting the geometric mean of housekeeping gene Ct values from the Ct value of each target gene to obtain ∆Ct; this value was transformed using the formula 2^-∆Ct^ to obtain a gene expression value for each subject (Livak and Schmittgen, 2001). Statistics were performed using Prism 6.0 (GraphPad Software, San Diego, CA) or Statistica (StatSoft Inc., Tulsa, OK). Outliers were determined by the robust regression and outlier analysis (ROUT) method with Q = 1% and were excluded from analyses (Motulsky and Brown, 2006). Two-tailed paired Student’s t-tests were performed for all genes unless 4 or more pairs were excluded after outlier removal; for the genes GNPTAB and ST8SIA6 primary statistical analysis was performed using unpaired Student’s t-tests. For the genes B3GNT3, GCNT3, and NEU2, which failed to amplify within 50 cycles (“No Ct” in instrument output) in 4 or more subjects, gene expression values were recorded as zero and the Wilcoxon matched pairs signed rank test was performed. We performed multiple testing correction using the Benjamini-Hochberg method (Benjamini and Hochberg, 1995)to control the false discovery rate with q* = 0.05. The experimental design employed a paired subject paradigm with no difference between groups for subject age, pH, or postmortem interval, however some of the genes under investigation in this study are known to exhibit age, sex, and/or medication related alterations in expression. In those cases, post hoc analyses of gene expression by demographic measure(s) for genes with altered expression in primary statistical tests were performed and are reported. For all tests, α = 0.05.

**Figure 1.**
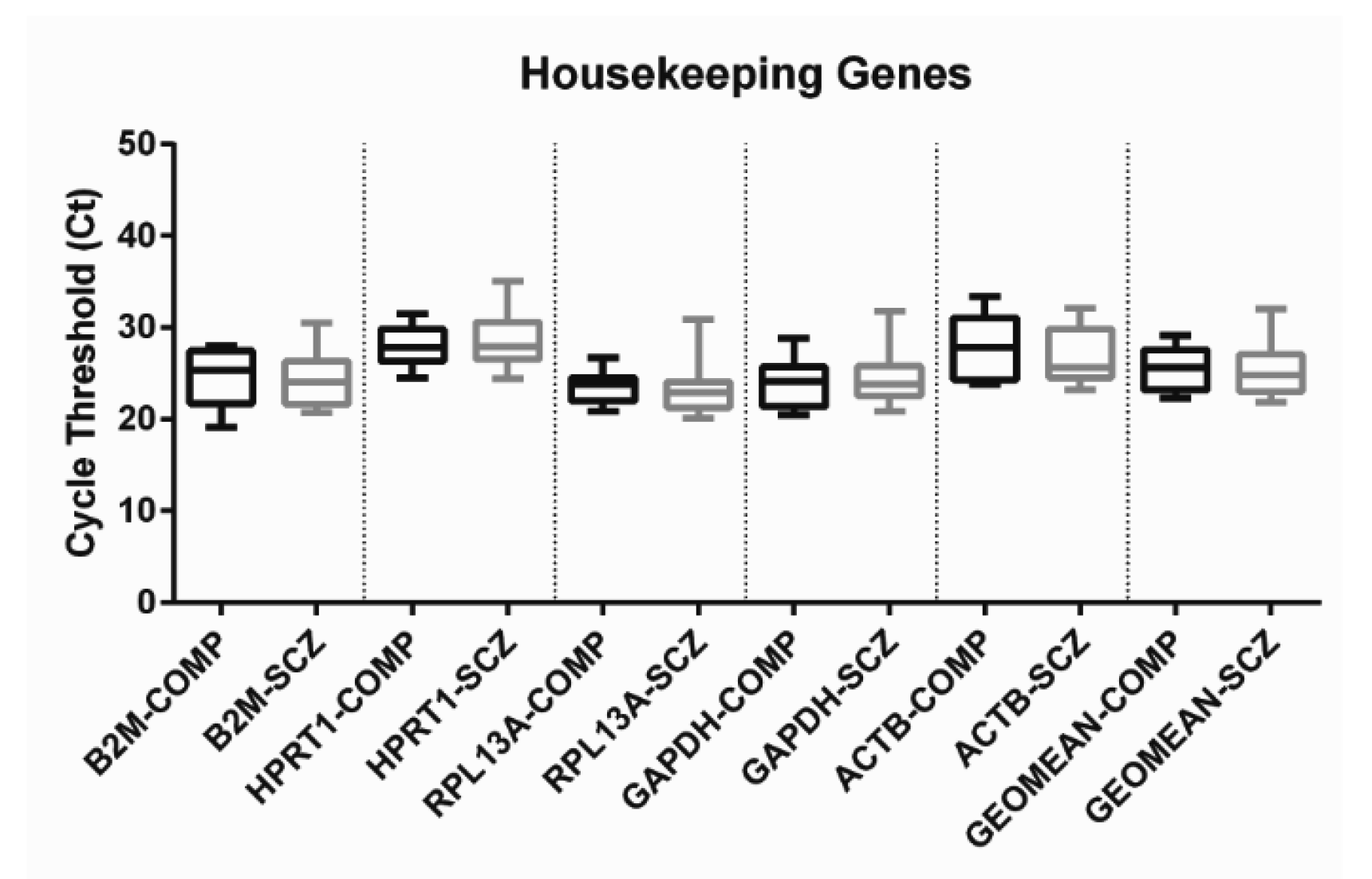
Housekeeping gene cycle threshold (Ct) values and the geometric mean of house-keeping gene Ct values are not different between diagnostic groups. Box-and-whisker diagrams of housekeeping gene cycle threshold (Ct) values and the geometric mean (geomean) of housekeeping genes. There is no difference between diagnostic groups of housekeeping gene Ct value for any of the individual housekeeping genes (B2M, HPRT1, RPL13A, GAPDH, and ACTB) nor the geomean of all housekeeping gene Ct values. The geomean of Ct values was used to calculate the gene expression of target glycosylation enzyme genes using the formula 2^-∆Ct^. To obtain ∆Ct, the geomean of housekeeping gene Ct values was subtracted from the target gene Ct value.

## Results

Glycosylation enzymes can be categorized based on whether they attach or remove glycans and the type of monosaccharide on which they act. In this study, enzymes from 16 functional categories were measured: fucosidases, fucosyltransferases, galactosidases, galactosyltransferases, glucosidases, glucosyltransferases, hexosaminidases, mannosidases, mannosyltransferases, N-acetylglucosaminidases, N-acetylglucosaminyltransferases, N-acetylgalactosaminidases, N-acetyl-galactosaminyltransferases, sialidases, sialyltransferases, and mannose-6-phosphate catabolic enzymes. Of the 84 glycosylation enzymes measured in this study, 46 genes were increased in schizophrenia relative to comparison subjects, and 36 remained significant after multiple testing correction using the Benjamini-Hochberg method (Table 3). Enzymes from all categories except galactosidases, hexosaminidases, mannosyltransferases, and mannose-6-phosphate catabolic enzymes had altered mRNA expression in schizophrenia. Enzymes that exert their primary activity in protein N-glycosylation, protein O-glycosylation, and sphingolipid processing pathways were found altered (Figure 2). None of the enzymes under investigation in this study had decreased transcript levels in schizophrenia.

**Table 3.**
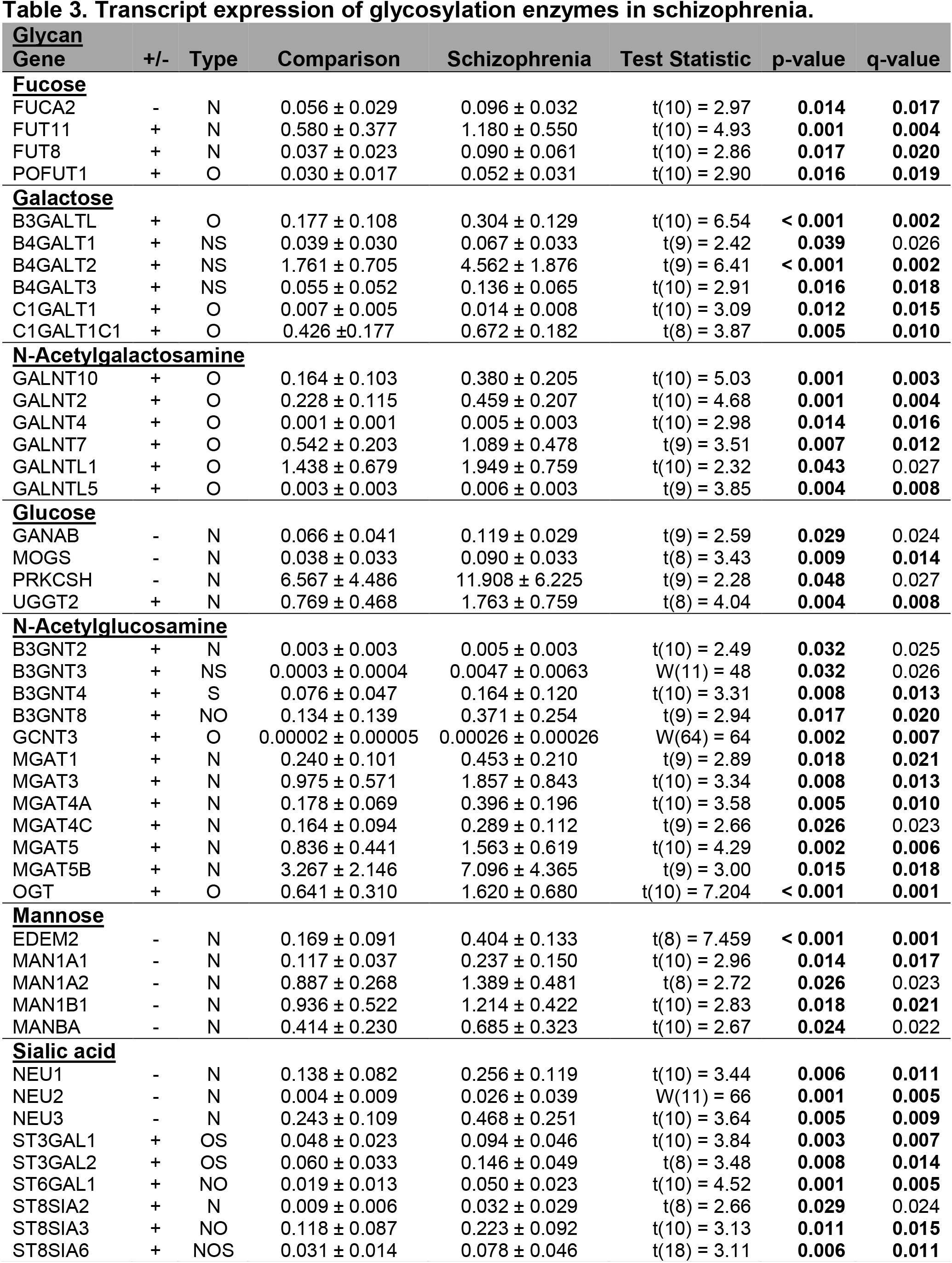
Transcript expression of glycosylation enzymes in schizophrenia.

**Figure 2.**
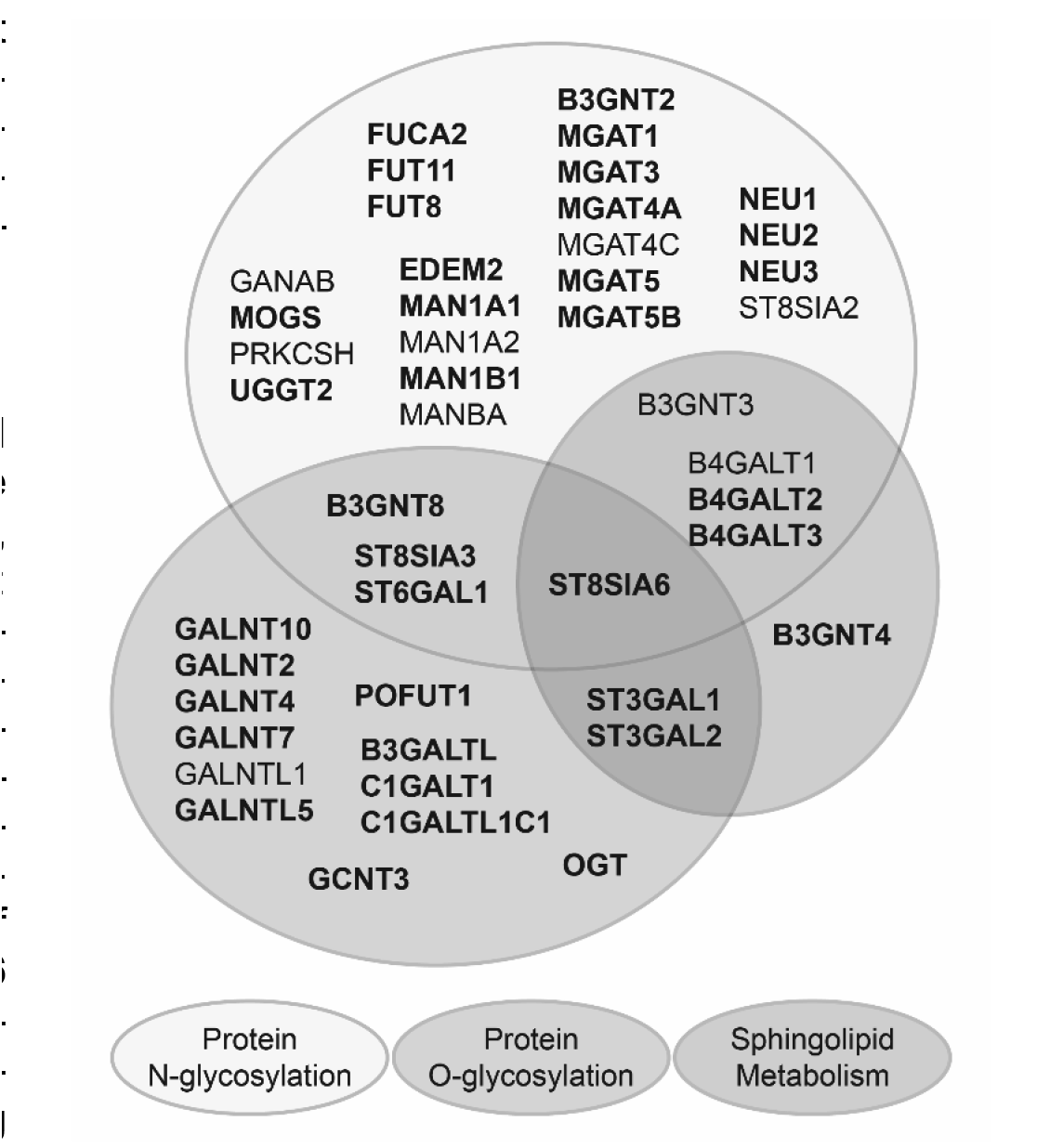
Summary of glycosylation enzyme transcript changes by pathway. Venn diagram of genes with significantly increased transcript expression in schizophrenia relative to paired comparison subjects (p < 0.05). Genes are grouped by the primary glycosylation pathway(s) in which they participate. Genes listed in bold survive multiple comparison correction (p < q, q* = 0.05).

### Enzymes Involved in Immature N-glycosylation in the ER are Altered in Schizophrenia

We identified increased mRNA levels of MOGS, MAN1A1, and MAN1B1, which play essential roles in N-glycan maturation and, with EDEM2 and UGGT2, which were also increased, contribute to ER quality control mechanisms (Figure 3) (Helenius and Aebi, 2004; Olivari and Molinari, 2007; Roth et al., 2010). Although we found no change in mRNA expression of MAN1C1, EDEM1, EDEM3, or UGGT1, and increased transcript levels of GANAB, PRKCSH, and MAN1A2 did not survive multiple testing correction, these data reflect dysregulated gene expression in schizophrenia of enzymes that participate in many stages of immature N-glycan processing (Figure 3).

**Figure 3.**
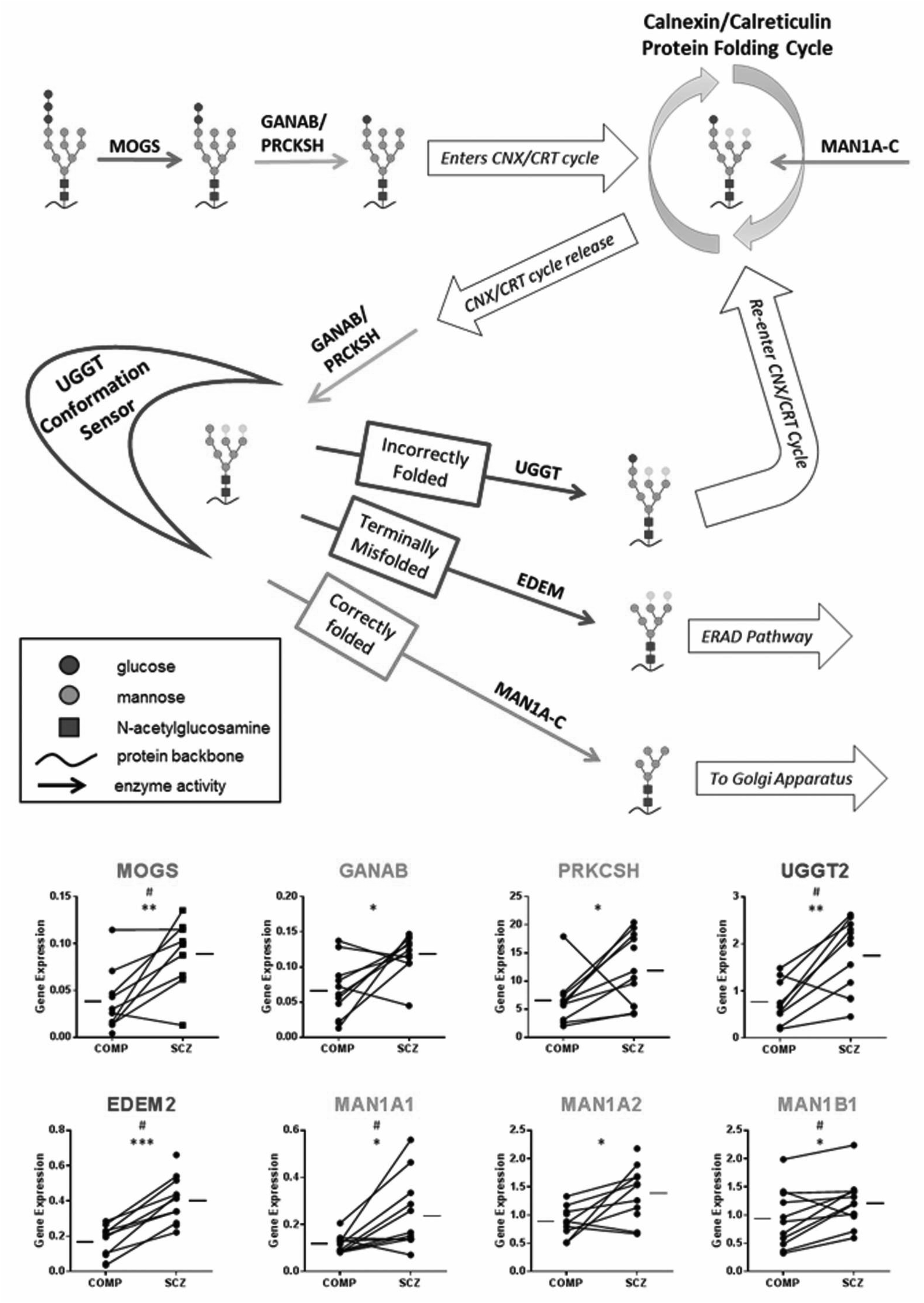
Transcript expression of immature N-glycosylation enzymes is increased in schizophrenia. Enzymes necessary for immature N-glycan processing have increased mRNA levels in schizophrenia. Top: Schematic of immature N-glycoprotein processing. After the N-glycan oligosaccharide precursor is attached to a protein substrate, α-glucosidase I (MOGS, red) cleaves the most terminal glucose. Cleavage of the next glucose by α-glucosidase II (GANAB, orange) is regulated by glucosidase II subunit β (PRCKSH, orange), and this N-glycan intermediate conformation allows the N-glycoprotein to interact with chaperone proteins in the calnexin-calreticulin (CNX/CRT) protein folding cycle. While interacting with molecules in the CNX/CRT cycle, ER-localized α-mannosidase I (MAN1A-C, green) slowly cleaves mannose residues. After N-glycoprotein release from the CNX/CRT protein folding cycle, GANAB/PRCKSH cleaves the final terminal glucose to allow interaction with UDP-glucose:glycoprotein glucosyltransferases (UGGT1 and 2, blue). UGGT proteins have two main functions: first, to act as conformation sensors to identify incompletely folded or misfolded proteins and, if the protein is not correctly folded, to next catalyze the re-addition of a terminal glucose to permit return to the CNX/CRT protein folding cycle. Correctly folded N-glycoproteins are released from the UGGT conformation sensor and can be further modified by MAN1A-C prior to forward trafficking to the Golgi compartment and further maturation into a complex N-glycan structure. If a protein has failed to attain the correct tertiary structure and is terminally misfolded, ER degradation-enhancing mannosidases (EDEM1-3, purple) interact with the glycoprotein after release from the UGGT conformation sensor and remove mannose residues to produce a Man^6^GlcNAc^2^-polypeptide structure which is exported from the ER and degraded via the ER-associated degradation pathway and ubiquitin proteasome system. Bottom: Graphs of gene expression of enzymes involved in immature N-glycan processing. Graph titles are color-coded to match the arrow color of enzymes in the diagram. Gene expression is calculated as 2^-∆Ct^ (arbitrary units). Data are from paired schizophrenia (SCZ) and comparison (COMP) subjects. *p < 0.05, **p < 0.01, ***p < .001, #p < q (survives Benjamini-Hochberg multiple testing correction). For statistical tests α = 0.05, q* = 0.05.

### Dysregulated Fucose-Modifying Enzyme Expression in Schizophrenia

Fucosyltransferases FUT11 and FUT8, protein O-fucosyltransferase POFUT1, and the fucosidase FUCA2 were increased in schizophrenia. POFUT2 and FUCA1 were not different between diagnostic groups. FUT8 has been shown to exhibit age-related increases in gene expression (Vanhooren et al., 2011) and, although we found no correlation between FUT8 expression and subject age in comparison subjects (r(11) = -0.24, p = 0.46), in schizophrenia there was a significant negative correlation between FUT8 expression and subject age (r(11) = -0.67, p = 0.02; Figure 4). To control for this effect, we performed an analysis of covariance (ANCOVA) of FUT8 gene expression by diagnostic group and subject age, and the difference between schizophrenia and comparison subjects remained significant (F(1,20) = 8.68, p = 0.008).

**Figure 4.**
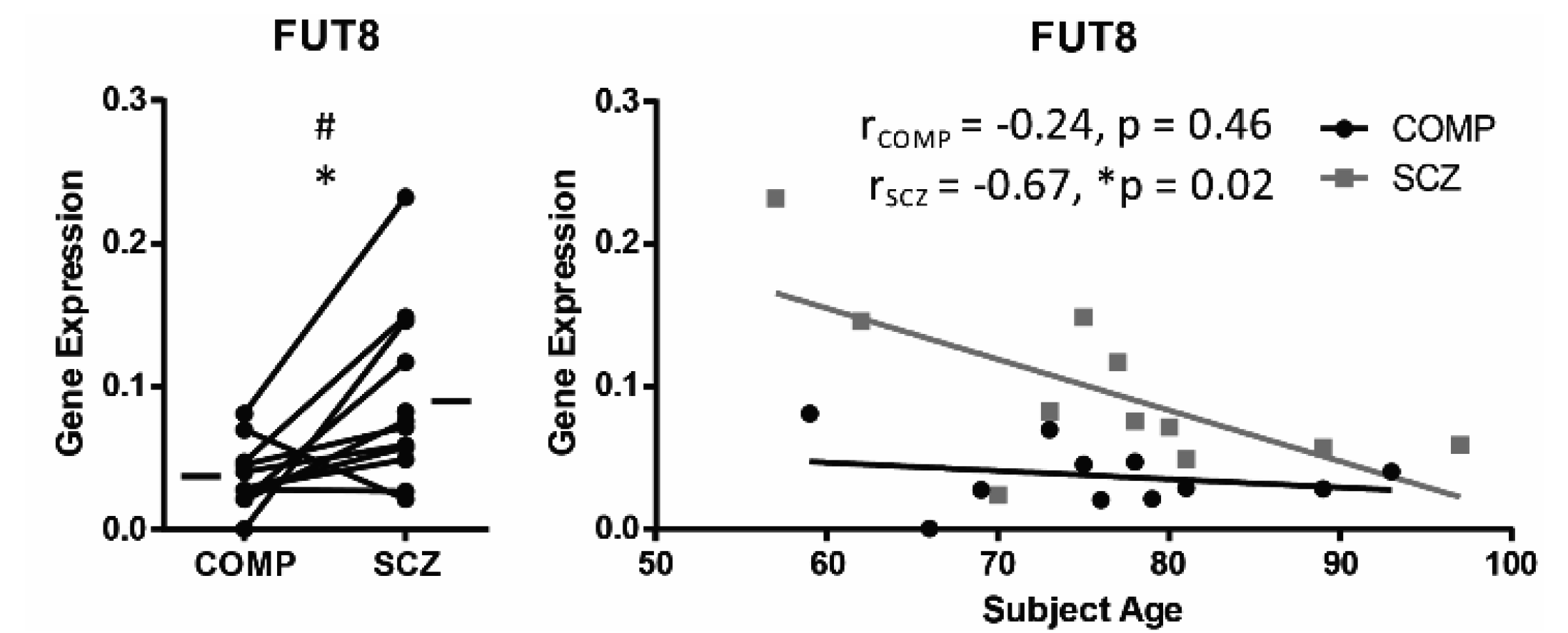
Increased FUT8 mRNA expression in schizophrenia inversely correlates with subject age. Graphs of gene expression of FUT8, calculated as 2^-∆Ct^ (arbitrary units). FUT8 mean gene expression is 44% higher in schizophrenia (SCZ) relative to comparison (COMP) subjects. There is no association between FUT8 gene expression and subject age in elderly COMP subjects; however, FUT8 expression declines with age in elderly SCZ subjects. Left: Data from paired SCZ and COMP subjects. Right: Gene expression correlated with subject age in SCZ and COMP subjects.*p< 0.05, #p < q (survives Benjamini-Hochberg multiple testing correction).For statistical tests α = 0.05, q* = 0.05.

### Altered N-acetylglucosaminyltransferase Expression in Schizophrenia

We assessed mRNA expression of multiple N-acetylglucosaminyltransferase (GlcNAcTase) subtypes: mannoside GlcNAcTases (MGATs), β-1,3-GlcNAcTases (B3GNTs), β-1,6-GlcNAcTases (B6GNTs), and the sole O-linked polypeptide GlcNAcTase. Transcripts encoding six MGATs (MGAT1, MGAT3, MGAT4, MGAT4B, MGAT5, and MGAT5B; Figure 5), two B3GNTs (B3GNT4 and B3GNT8), one B6GNT (GCNT3) and O-GlcNAcTase (OGT) were increased in schizophrenia. There was no difference in expression of MGAT2, MGAT4B, MGAT4C, B3GNT2, B3GNT3 GCNT1, and GCNT4 between diagnostic groups.

**Figure 5.**
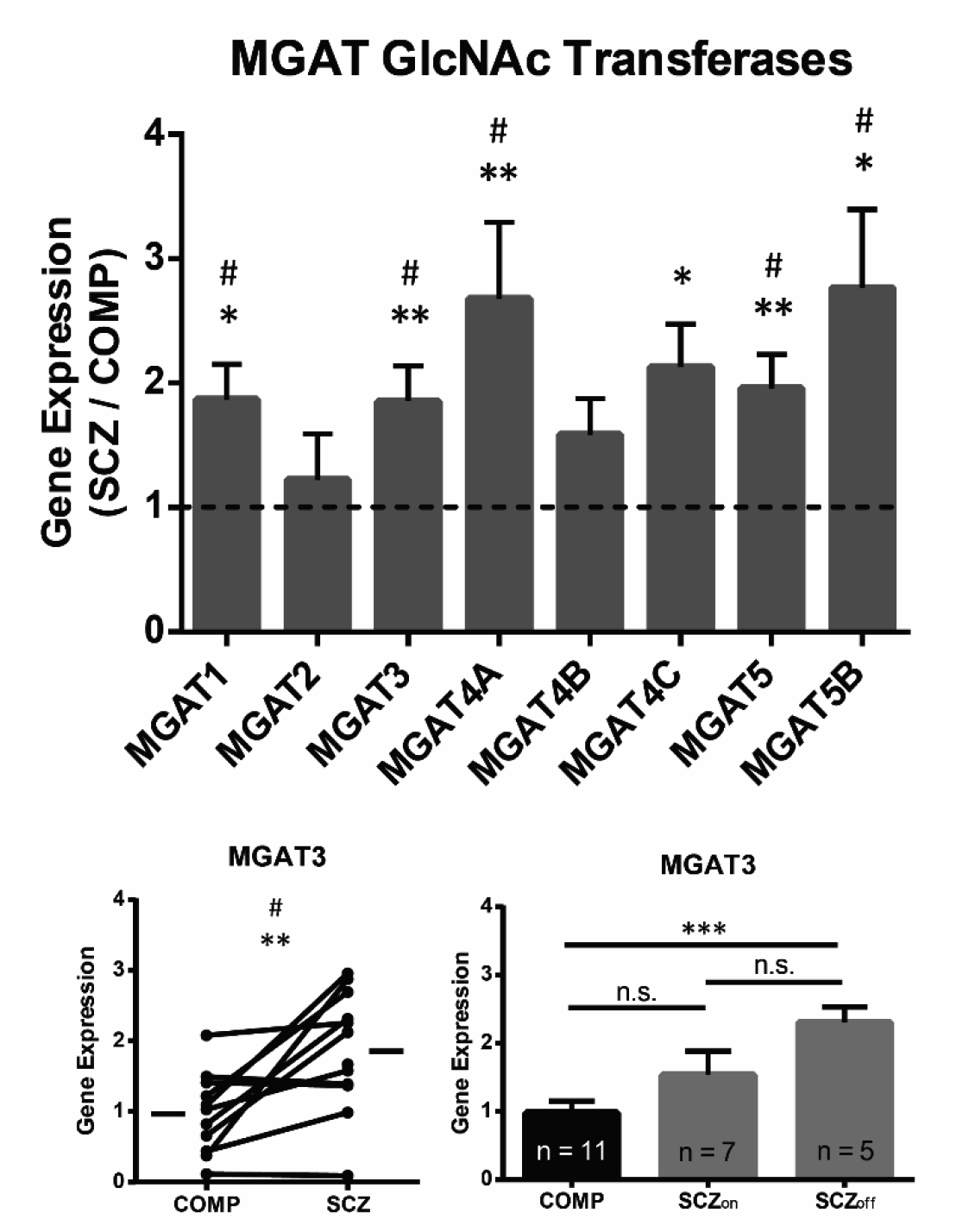
Transcript levels of MGAT1, MGAT3, MGAT4A, MGAT5, and MGAT5B are increased in schizophrenia. Gene expression of MGAT enzymes calculated as 2^-∆Ct^ (arbitrary units). Gene expression of MGAT1, MGAT3, MGAT4A, MGAT5, and MGAT5C is increased in schizophrenia (SCZ) relative to comparison (COMP) subjects. The difference in gene expression of MGAT3 between diagnostic groups is driven by increased expression in SCZ subjects who had not received antipsychotic medication within 6 weeks of death (SCZ_off_), and the mRNA levels of MGAT3 in SCZ subjects on antipsychotic medication (SCZ_on_) is intermediate between SCZ_off_ and COMP subjects. Top: Gene expression in SCZ divided by gene expression in matched COMP subject, data are expressed as means ± S.E.M. Bottom left: MGAT3 gene expression in matched SCZ and COMP subjects. Bottom right: Post hoc analysis of MGAT3 gene expression between COMP, SCZon, and SCZ_off_. Data are represented as means ± S.E.M. *p< 0.05, **p < 0.01, ***p < .001, #p < q (survives Ben-jamini-Hochberg multiple testing correction), n.s. = not significant. For statistical tests α = 0.05, q* = 0.05.

Lower gene expression of MGAT3 in the DLPFC of schizophrenia subjects within 5 years of diagnosis and a reduction in biantennary bisected glycans (suggesting decreased MGAT3 activity) in CSF from antipsychotic naïve schizophrenia patients have previously been reported; however, these measures were obtained during the early stages of illness progression in younger adults (Narayan et al., 2009; Stanta et al., 2010). Given our finding of increased MGAT3 mRNA expression, we performed post hoc analyses to address whether increased MGAT3 expression may be an effect of advanced age or the medication status of the subjects used in this study. We performed regression analysis of MGAT3 gene expression by subject age and found no significant correlation. Analysis of MGAT3 gene expression between comparison subjects, schizophrenia subjects on medication (SCZ_on_), and schizophrenia subjects off medication (SCZ_off_; defined as subjects that did not receive antipsychotics for 6 weeks or more at time of death) revealed no difference in expression between SCZ_on_ and SCZ_off_ subjects; however, the difference between comparison and schizophrenia subjects is primarily driven by increased expression in SCZ_off_ subjects (Figure 5).

It is possible that B3GNT3 and GCNT3 are not normally expressed at detectible levels in human DLPFC of elderly subjects (Figure 6). These enzymes failed to meet the threshold for detection within 50 qPCR cycles in at least 3 comparison subjects. Mean gene expression of B3GNT3 and GCNT3 were an order of magnitude higher in schizophrenia relative to comparison subjects.

**Figure 6.**
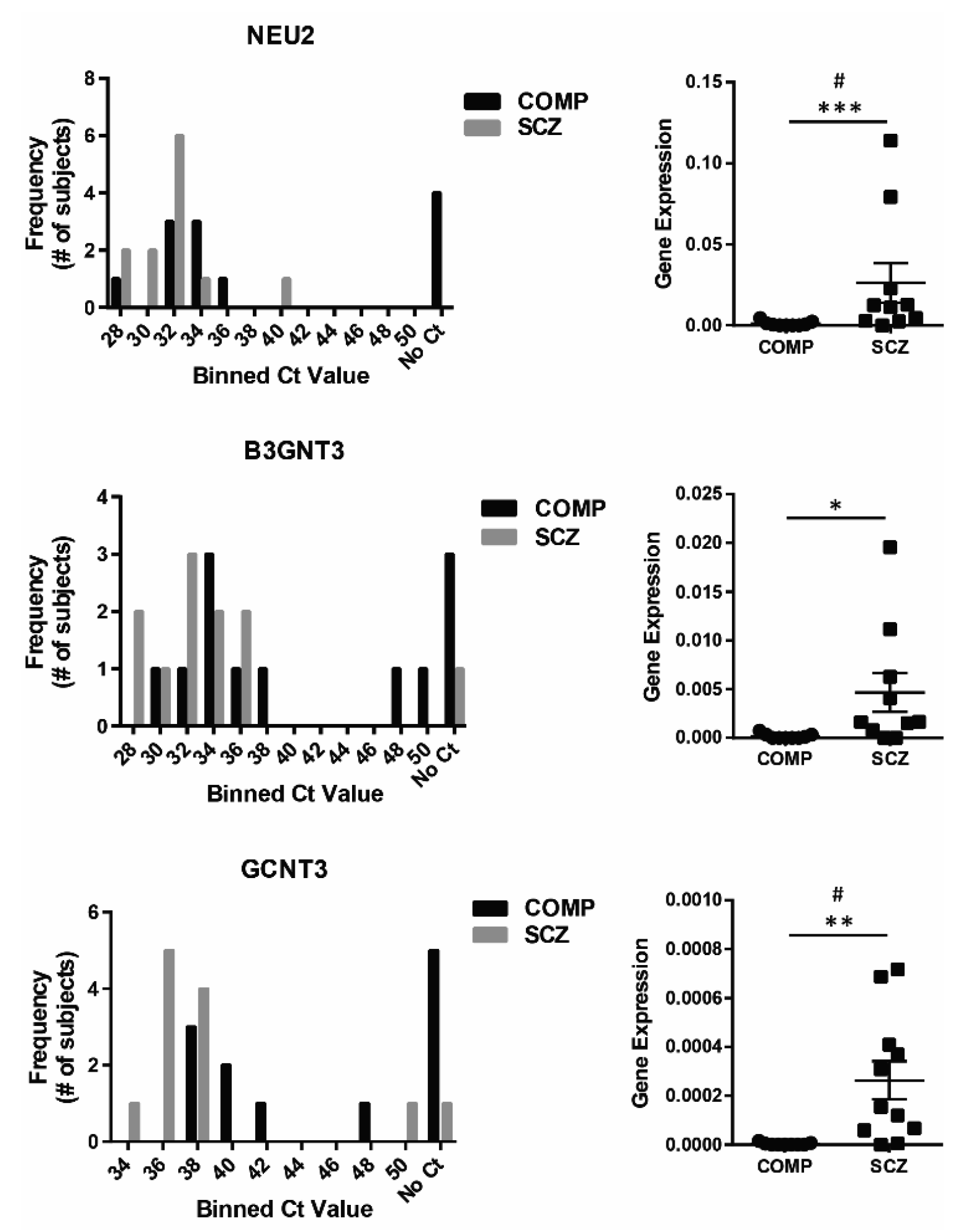
NEU2, B3GNT3, and GCNT3 mRNA expression in postmortem cortex of elderly subjects. Subject mRNA was reverse transcribed to cDNA and samples of cDNA were preamplified prior to qPCR to permit the assessment of low abundance transcripts in postmortem human brain. In spite of this preamplification step, some enzymes--NEU2, B3GNT3, and GCNT3—failed to amplify within 50 qPCR cycles in some subjects. These enzymes were detected at levels at least an order of magnitude higher in SCZ (SCZ) than (COMP) subjects and may not normally be expressed at detectable levels in neurotypical cortex. Gene expression is calculated as 2^-∆Ct^ (arbitrary units); for subjects that failed to amplify, gene expression value was recorded as zero. Left: Histograms of raw cycle thresh-old (Ct) values; bin size is 2 cycles. Right: Graphs of individual subjects with means ± S.E.M. indicated, *p< 0.05, **p < 0.01, ***p < .001, #p < q (survives Benjamini-Hochberg multiple testing correction).

### Sialyltransferase and Sialidase Expression Abnormalities in Schizophrenia

We found increased mRNA expression of five sialyltransferases in schizophrenia: ST3GAL1, ST3GAL2, ST6GAL1, ST8SIA3, and ST8SIA6. Although deficits associated with mutations of ST8SIA2 have been implicated in the decreased expression of PSA-NCAM in schizophrenia cortex (Barbeau et al., 1995; Isomura et al., 2011), ST8SIA2 gene expression was not decreased in this study. Transcript expression of the sialidases NEU1, NEU2, and NEU3, but not NEU4, were increased in schizophrenia. In comparison subject DLPFC, NEU2 gene expression was approximately 2 orders of magnitude less than NEU1 and NEU3 expression; however, while NEU1 and NEU3 expression was increased approximately 86% and 92%, respectively, in schizophrenia, NEU2 expression was increased over 600% from comparison levels. NEU2 also failed to meet the threshold for detection within 50 qPCR cycles in four comparison subjects, but demonstrated amplification within 40 cycles in all of the schizophrenia subjects (Figure 6).

### Altered Galactose and N-Acetylgalactosamine Associated Enzymes in Schizophrenia

Transcripts encoding four enzymes from the galactosyltransferase family were found increased in schizophrenia: C1GALT1, B4GALT2, B4GALT3, and B3GALTL. C1GALT1C1 (also called COSMC), which is a chaperone for C1GALT1, was also increased. N-acetylgalactosaminyltransferases (GalNAcTases) GALNT2, GALNT4, and GALNT10, which initiate protein O-GalNAcylation, and GALNT7 were increased in schizophrenia (Bennett et al., 1999, 1998; Fritz et al., 2006; Moremen and Nairn, 2014). Other galactosyl-transferases and GalNAcTases measured in this study were not altered in schizophrenia.

## Discussion

These data demonstrate that genes involved in multiple glycosylation pathways are abnormally expressed in postmortem schizophrenia DLPFC. Previous findings from our lab have demonstrated in schizophrenia abnormalities of N-linked glycosylation of neurotransmitter receptor and transporter subunits in multiple brain regions including the DLPFC, anterior cingulate cortex, and superior temporal gyrus (Bauer et al., 2010; Mueller et al., 2014; Tucholski et al., 2013a, 2013b). It has also been shown that the glycan profile in serum and CSF of first onset, antipsychotic naïve schizophrenia patients is distinct from that of comparison subjects (Stanta et al., 2010) and that treatment with olanzapine results in abnormal glycan branching of serum glycoproteins (Telford et al., 2012). Together, these data represent a growing body of evidence implicating disruptions in glycan processing in schizophrenia pathophysiology. To reconcile these diverse findings, we examined the transcript expression of 84 enzymes involved in N-glycosylation, O-glycosylation, sphingolipid metabolism, and/or mannose-6-phosphate catabolism in DLPFC. In schizophrenia, 46 genes demonstrated increased expression relative to comparison subjects and 36 of these genes survived multiple testing correction (Table 3).

The majority of altered transcripts were for enzymes that mediate glycan attachment (glycosyltransferases), whereas only 7 of the altered enzymes are responsible for glycan cleavage (glycosidases). We predicted changes in ER-localized glycosylation enzymes in schizophrenia, and found 8 altered enzymes that are specifically expressed in the ER (Figure 7). Given our prediction of dysfunction in multiple glycosylation pathways, it is noteworthy that the majority of abnormally expressed enzyme transcripts are primarily expressed in the Golgi and/or the extracellular space (Binder et al., 2014), locations where glycosylation pathways converge and glycan modifications are of particular functional importance (Varki et al., 2009). Of the 36 genes with increased transcript expression in schizophrenia, 16 are exclusively associated with N-glycosylation, 11 are exclusively associated with O-glycosylation, 1 is exclusively associated with sphingolipid metabolism, and 8 are involved in multiple protein glycosylation pathways (Table 3, Figure 2).

**Figure 7.**
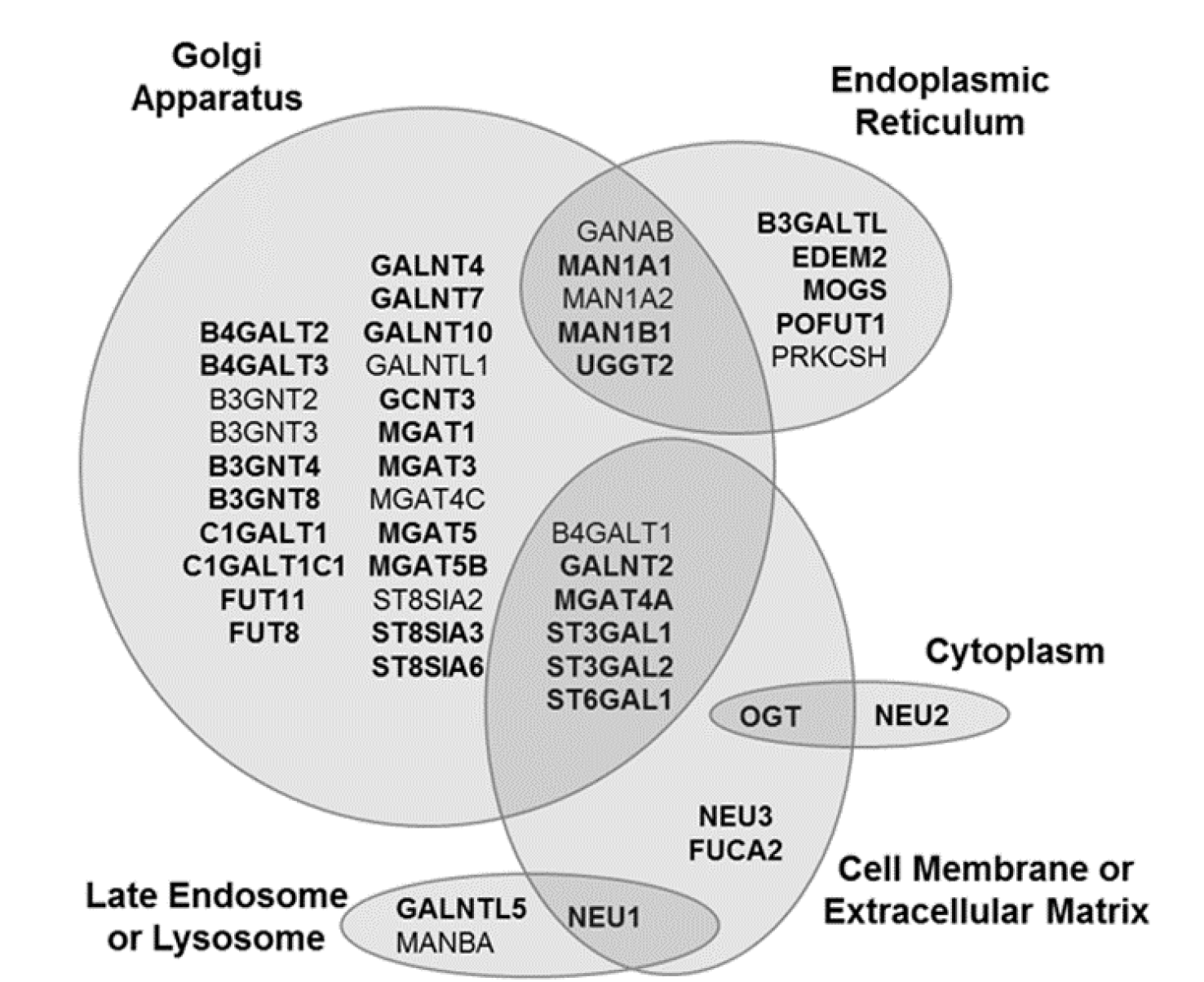
Subcellular distribution of glycosylation enzymes found altered in schizophrenia. Venn diagram of genes with significantly increased transcript expression in schizophrenia relative to paired comparison subjects (p < 0.05). Genes are grouped by the primary subcellular location(s) of the gene product between the endoplasmic reticulum, Golgi apparatus, cytoplasm, late endosome or lysosome, and cell membrane or extracellular matrix based on assignments curated by and reported in the UniProtKB (http://www.uniprot.org) and COMPARTMENTS (http://compartments.jensenlab.org) databases. The “cell membrane or extracellular matrix” category includes enzymes that associate with the cytoplasmic side of the plasma membrane, transmembrane proteins expressed on the cell surface, secreted proteins that interact directly with the proteinaceous extracellular matrix, or proteins secreted via extracellular exosomes. The enzyme OGT is functionally active in cell nuclei and within mitochondria, as well as throughout the cytoplasm and at the cell membrane as shown. Some glycosylation enzymes may also localize to other specific subcellular compartments not indicated here. Genes listed in bold survive multiple comparison correction (p < q, q* = 0.05).

### Immature N-glycosylation in the ER and Glycan Branching

The co- or post-translational attachment of an oligosaccharide precursor to an asparagine residue initiates N-linked glycosylation of proteins in the ER and subsequent glycan processing in a specific, step-wise manner acts as an intracellular signal to identify abnormally folded proteins (Figure 2, top) (Helenius and Aebi, 2004; Roth et al., 2010; Varki et al., 2009). Increased transcription of MOGS, MAN1A1, MAN1B1, EDEM2, and UGGT2, were identified in schizophrenia. Given our prior reports of abnormal immature N-glycosylation of the α1 and β1 GABA_A_ receptor subunits (Mueller et al., 2014), the GRIA2 AMPA receptor subunit (Tucholski et al., 2013a), and the GRIK2 kainate receptor subunit (Tucholski et al., 2013b), it is not surprising to find abnormal expression of these enzymes. These data support our hypothesis that abnormal immature N-glycosylation in schizophrenia arises from abnormal expression of glycosylation enzymes and increased transcription of MOGS, MAN1A1, MAN1B1, UGGT2, and EDEM2 may therefore represent an underlying mechanism of disrupted immature N-glycoprotein processing and quality control functions in the ER in schizophrenia. Preliminary data from our lab indicates that concurrent with increased mRNA levels, protein expression of EDEM2 and UGGT2 are increased in schizophrenia DLPFC (Kim et al., 2018). Together with evidence of dysregulated degradation pathways in schizophrenia (Kim et al., 2018; Rubio et al., 2013; Scott et al., 2015), these findings suggest that upregulation of ER-resident glycan modifying enzymes may alter the basal activity of the ER-associated degradation (ERAD) and unfolded protein response (UPR) pathways in schizophrenia.

When an N-glycoprotein has attained the correct conformation in the ER, the high-mannose N-glycoprotein is exported to the Golgi apparatus and can be modified by further cleavage of mannose residues and/or attachment of additional monosaccharides (Varki et al., 2009). The MGAT family of GlcNAcTases converts immature, high-mannose N-glycans to hybrid or complex-type N-glycans by the addition of GlcNAc to the N-glycan trimannosyl core (Kumar et al., 1990; Schachter, 1986; Yip et al., 1997). Sequential activity of MGAT1 and 2 in the Golgi precedes the competition of MGAT3, 4, and 5 to produce bi-, tri-, and tetra-antennary glycan structures, respectively, and altered expression of any of these enzymes can affect final N-glycan conformations significantly for a wide variety of substrates (Kippe et al., 2015; Moremen and Nairn, 2014; Ohtsubo and Marth, 2006; Schachter, 1986). Expression levels of mRNA for MGAT1, MGAT3, MGAT4, MGAT5, and MGAT5B were increased in schizophrenia. A concurrent study in our lab found the protein expression of MGAT4A to be significantly decreased in schizophrenia, and we have proposed that the opposite valence of mRNA and protein findings for this enzyme likely represents a compensatory mechanism in the disorder (Kippe et al., 2015). To reconcile our findings with prior reports of decreased MGAT3 mRNA in younger antipsychotic-naïve schizophrenia subjects, post hoc analyses of effects of age and antipsychotic treatment on MGAT3 gene expression were performed. We identified no significant correlation with age in elderly subjects. There was also no difference of gene expression found between SCZ_on_ and comparison or SCZ_off_ subjects, but transcript expression of MGAT3 in SCZ_off_ was increased relative to comparison subjects (Figure 3). These findings suggest that increased gene expression of MGAT3 may occur at a later stage of the disorder rather than as an effect of aging or antipsychotic treatment. Additionally, given that the MGAT3 mRNA expression level in SCZ_on_ subjects was intermediate between SCZ_off_ and comparison subjects, one might speculate that antipsychotic treatment may ameliorate some glycosylation-associated abnormalities in schizophrenia.

### Protein O-GlcNAcylation

A specific type of O-glycosylation, protein O-GlcNAcylation, is of particular interest due to the cyclical nature of O-GlcNAcylation and phosphorylation on shared serine/threonine residues (Moremen and Nairn, 2014). Unlike other forms of protein N- or O-glycosylation which occur in the ER or Golgi, protein O-GlcNAcylation is initiated in the nucleus or cytosol (Ryšlavá et al., 2013). The phosphorylation/O-GlcNAcylation cycle acts as a substrate-specific intracellular signal which can regulate gene transcription, cytoskeletal remodeling, and nutrient homeostasis, and is initiated by OGT (Butkinaree et al., 2010; Howerton et al., 2013), which we found increased in schizophrenia. Given the potential competition of O-GlcNAcylation and phosphorylation on an estimated 30% of mammalian glycoproteins (Ryšlavá et al., 2013) and abnormal kinase, phosphatase, and phosphoprotein expression in schizophrenia (Kim and Stahl, 2010; McGuire et al., 2017, 2014), alterations of O-GlcNAcylation could potentially contribute to altered cellular function in schizophrenia. The OGT gene is located on the X-chromosome, and expression of this enzyme is lower in males than females in neurodevelopment (Howerton et al., 2013). Recently, abnormal OGT expression has been correlated with sex-biased neurodevelopmental alterations following maternal stress exposure (Howerton et al., 2013). Schizophrenia demonstrates sex-specific variations in illness onset, progression, and pathophysiology (Häfner et al., 1994), and epidemiological findings have correlated maternal stress with increased schizophrenia risk (Bayer et al., 1999; Khandaker et al., 2013). Together these data suggest a possible functional effect of abnormal OGT expression in sex-specific features of schizophrenia.

### Sialylation and Polysialylation of Membrane Proteins

The attachment of N-acetylneuramic acid (known commonly as sialic acid and abbreviated NeuAc) onto membrane proteins and glycolipids is a modification which influences the strength and stability of synaptic connections and cell-cell interactions (Dennis et al., 2009; Maccioni et al., 2011; Varki et al., 2009). Many glycans, particularly on cell surface glycoproteins and gangliosides, are capped by the addition of NeuAc or poly-NeuAc which prevents further monosaccharide additions (Maccioni et al., 2011; Schnaar et al., 2014). Reports of abnormal protein sialylation and reduced levels of the polysialylated form of NCAM (Barbeau et al., 1995; Isomura et al., 2011; Stanta et al., 2010; Telford et al., 2012) are implicated in abnormal cell-cell interactions, atypical neurodevelopment, and synaptic abnormalities in schizophrenia (Barbeau et al., 1995; Schnaar et al., 2014). We found mRNA of α-2,8-sialyltransferases ST8SIA3 and ST8SIA6, α-2,3-sialyltransferases ST3GAL1 and ST3GAL2, and α-2,6-sialyltransferase ST6GAL1 increased in schizophrenia. It has previously been reported that ST3GAL2 is not expressed in human brain based on northern blot analysis (Giordanengo et al., 1997), however, the prior study did not specify which brain region was assessed for ST3GAL2 gene expression and our current study measured expression using a different methodology.

### Synthesis of Lactose, N-Acetyllactosamine, and Poly-N-Acetyllactosamine

B4GALT2 catalyzes the synthesis of lactose or N-acetyllactosamine (LacNAc) from UDP-galactose and glucose or GlcNAc, respectively, and adds glucose to glycolipids or to N-linked oligosaccharides to produce complex N-glycoproteins (Guo et al., 2001). B4GALT3 modifies glycolipids and N-glycoproteins as well as playing a role in the synthesis of LacNAc, but not lactose (The UniProt Consortium, 2014). Similar to the MGAT family of GlcNAcTases, which are involved in N-glycan maturation, the B3GNTs also extend existing glycans and synthesize specific branched glycan structures (Moremen and Nairn, 2014; Varki et al., 2009). B3GNT4 modifies glycolipids and is involved in the generation of poly-LacNAc, primarily in microglia, ependymal cells, and brain vasculature (Acarin et al., 1994; Shiraishi et al., 2001; The UniProt Consortium, 2014). B3GNT8 can also synthesize poly-LacNAc but cannot modify sphingolipids (Ishida et al., 2005; Seko and Yamashita, 2005). Similar to the GlcNAcTase MGAT4A, discussed in section 4.1, protein expression of B3GNT8 was also found to be decreased in schizophrenia DLPFC, and increased mRNA levels reported here may reflect cellular compensation for inadequate protein levels of the enzyme (Kippe et al., 2015). Increased gene expression of these enzymes in our elderly cohort is consistent with previous reports of altered expression of enzymes involved in the synthesis of lactoseries glycosphingolipids in schizophrenia (Narayan et al., 2009). Lactoseries glycosphingolipids are developmentally regulated membrane-bound glycolipids that exhibit two main cellular functions: regulation of cell-cell interactions via contact with molecules on the apposing cell membrane and functional modulation of proteins expressed in the same cell membrane (Varki et al., 2009; Yu et al., 2011). Interestingly, abnormal sphingolipid expression can produce deficits of attention and impulse control, and glycosphingolipid-mediated regulation has been suggested to play a role in maintaining neuropsychological balance (Kuan et al., 2010; Niimi et al., 2011; Yu et al., 2011).

### Extracellular Matrix and Synaptic Remodeling

C1GALT1 and C1GALT1C1 (also called COSMC) must be co-expressed to generate the T antigen necessary for core 1 O-glycosylation, and C1GALT1C1 expression is a limiting factor in the synthesis of core 1 O-glycans, so it is unsurprising to find C1GALT1 and C1GALT1C1 increased concurrently in schizophrenia. Many secreted glycoproteins are extensively O-glycosylated (Varki et al., 2009). B3GALTL in the ER is responsible for elongating O-fucosylproteins following POFUT2-mediated fucosylation of thrombospondin-like repeat (TSR) domains (Sato et al., 2006) and for some ADAMTS (<u>a</u> <u>d</u>isintegrin <u>a</u>nd <u>m</u>etalloproteinase with thrombo<u>s</u>pondin motifs) proteins, the O-linked disaccharide (produced from sequential glycan addition by POFUT2 and B3GALTL) has been shown to be necessary for secretion into the extracellular space (Ricketts et al., 2007; Wang et al., 2007). We have recently reported that the protein expression levels of POFUT2 in the superior temporal gyrus (STG; Brodmann area 22) is decreased in schizophrenia (Mueller et al., 2017). While we did not identify mRNA expression changes of POFUT2 in the current study, this may reflect a brain-region specific alteration in POFUT2 protein levels or the increased transcript levels of B3GALTL identified in DLPFC may arise from inadequate or abnormal TSR-domain O-fucosylation. Secreted ADAMTS proteins cleave lecticans to permit synaptic remodeling (Gottschall and Howell, 2015; Ricketts et al., 2007; Sato et al., 2006; Wang et al., 2007) and abnormal expression of either or both B3GALTL and POFUT2 could contribute to alterations of synaptic plasticity in schizophrenia by impacting ECM remodeling in the disorder.

POFUT1 is responsible for protein O-fucosylation and specifically modifies epidermal growth factor homology (EGF) domains when proper disulfide pairing of cysteines has occurred (Harris and Spellman, 1993; Wang et al., 2001). An important family of POFUT1 substrates are the NOTCH proteins, which are known for their role in cell fate determination and neurogenesis during development and which influence neuritogenesis and dendritic spine formation in post-mitotic adult neurons (Berezovska et al., 1999; Lai, 2004; Redmond et al., 2000). Appropriate NOTCH protein O-glycosylation is necessary for both the correct tertiary structure of the glycoprotein and for the conformation shift that occurs following ligand binding (Irvine, 2008). Alterations of the NOTCH signaling pathway have been implicated in schizophrenia (Pietersen et al., 2014; Shayevitz et al., 2012; Yang et al., 2013), and these data suggest abnormal NOTCH fucosylation may contribute to disruption of this signaling pathway.

FUT8 and FUT11 are N-linked fucosyltransferases increased in schizophrenia. FUT11 is one of the most evolutionarily conserved α-1,3-fucosyltransferases and there is strong sequence similarity between FUT11 and other α-1,3-fucosyltransferases (Baboval and Smith, 2002; Mollicone et al., 2009). Conversely, FUT8 is unique in its ability to catalyze the addition of α-1,6-linked fucose onto the chitobiose core of N-linked glycans (Kötzler et al., 2012). FUT8 knock-out mice demonstrate a schizophrenia-like behavioral phenotype, and many neurodevelopmental pathways that regulate neuritogenesis and synaptic strengthening that have been implicated in schizophrenia are dysregulated when N-glycoproteins are not appropriately core fucosylated (Fukuda et al., 2011; Gu et al., 2015, 2013; Kurimoto et al., 2014; Wang et al., 2006, 2005). Previous studies have reported FUT8 expression increases in normal aging in mice (Vanhooren et al., 2011), and while we did not find a correlation between FUT8 expression and subject age in human comparison subjects, we did find a negative correlation in schizophrenia. This may suggest abnormalities of brain aging in schizophrenia that are consistent with atypical neurodevelopment in earlier stages of the disorder. As with the fucosyltransferase POFUT2, protein levels of FUT8 were found decreased in schizophrenia STG (Mueller et al., 2017), supporting the notion that glycan processing enzyme expression protein and/or mRNA levels demonstrate brain region-specific abnormalities in schizophrenia.

### Degradation Pathways

We found increased mRNA expression of the sialidases NEU1, NEU2, and NEU3, which are expressed in the lysosome, cytosol, and plasma membrane, respectively (Albohy et al., 2010; Binder et al., 2014; Miyagi et al., 2008). NEU1-4 are the only known sialidases expressed in humans and although they all cleave NeuAc, substrate specificity and enzymatic properties are unique for each enzyme (Magesh et al., 2006; The UniProt Consortium, 2014). NEU1 acts primarily within the lysosomal compartment and has been shown to negatively regulate lysosomal exocytosis, an autophagic process (Bonten et al., 1996; d’Azzo and Bonten, 2010; Yogalingam et al., 2008). Conversely, NEU3 inhibits matrix metalloproteinase 9 (MMP9) expression (Moon et al., 2007) and can increase intracellular lactosyl ceramide to induce beclin-2 expression, thereby inhibiting apoptosis and enhancing cell survival and autophagy pathways (Kato et al., 2006; Magesh et al., 2006; Scaringi et al., 2013; Tringali et al., 2009). MMP9 inhibition also leads to decreased NEU1 activity, proposed to be a result of increased NEU3 expression, and can lead to hyperglycemia and insulin resistance (Alghamdi et al., 2014), a side-effect of many antipsychotic drugs (Heal et al., 2012).

Deficits of GALNTL5 expression have been shown to result in abnormal localization and functional deficits of the ubiquitin-proteasome system (UPS) in sperm cells (Takasaki et al., 2014; The UniProt Consortium, 2014). In schizophrenia brain, we found this enzyme upregulated. While specific consequences of this increase are unclear, abnormal expression of proteins involved in the UPS in schizophrenia brain have recently been reported (Rubio et al., 2013; Scott et al., 2015), and cross-talk between glycosylation and ubiquitination pathways in schizophrenia may suggest a possible mechanism underlying altered UPS function in the disorder.

The pH-dependent lysosomal and extracellular fucosidase FUCA2 can be upregulated in response to immune activation (Sobkowicz et al., 2014), and we found increased mRNA expression of FUCA2 in schizophrenia. The substrates and role of FUCA2 in brain have not been well-characterized, though as we have identified increased transcript levels of fucosyltransferases in schizophrenia, it is reasonable to expect that this increase may be a mechanism by which neurons attempt to compensate for increased fucosylation of some substrates. Taken together, the abnormal expression of lysosomal fucosidase (FUCA2), lysosomal sialidase (NEU1), ERAD-associated mannosidase (EDEM2; discussed in section 4.1), and UPS-associated (GALNTL5) glycosylation enzymes suggest atypical function of multiple protein degradation pathways in schizophrenia and may represent novel targets for future investigation.

### Limitations and Conclusions

A common limitation of studies in postmortem schizophrenia brain is the potential confound of patient medication status at time of death. We have defined 6 weeks of antipsychotic abstinence prior to death as “off” medication; however, given the advanced age of schizophrenia subjects included in this study, treatment with antipsychotic medication occurred at some point during the patients’ life and 6 weeks may not be sufficient time to reverse alterations arising from previous chronic antipsychotic use. Given the dynamic regulation of glycosylation enzyme expression, additional studies in antipsychotic-naïve subjects and subjects in earlier stages of the disorder are warranted.

We have speculated on potential consequences of abnormal gene expression of glycosylation enzymes in schizophrenia; however, inconsistencies between mRNA abundance and protein levels of the corresponding gene product from samples of postmortem human brain are not uncommon (Harrison and Weinberger, 2005; Kristiansen et al., 2007; Meador-Woodruff and Healy, 2000). We have recently reported decreased protein expression of the GlcNAcTases B3GNT8 and MGAT4A in the DLPFC of these same subjects (Kippe et al., 2015), and decreased protein expression of the fucosyltransferases FUT8 and POFUT2 in the STG in a different subject cohort (Mueller et al., 2017), while both mRNA and protein levels of EDEM2 and UGGT2 are increased in schizophrenia DLPFC. The data presented herein do not contradict our prior reports, but instead suggest that increased mRNA expression may be cellular compensation for insufficient protein expression of these enzymes in schizophrenia. We consider this finding support for our hypothesis that abnormal glycosylation in schizophrenia may be related to subcellular compartment-specific abnormalities in glycan processing. B3GNT8, MGAT4A, and FUT8 are primarily localized to the Golgi apparatus, thus the opposite valence of protein and mRNA expression changes may be an indication that glycosylation alterations in schizophrenia stem from abnormalities of glycan or glycoprotein processing in the ER which the cell attempts to compensate for further along the processing pathway in the Golgi apparatus. While the expression of some other GlcNAcTases found to be abnormally transcribed in this study were found unchanged at the protein level (Kim et al., 2018; Kippe et al., 2015; Mueller et al., 2017), this does not preclude the possibility that PTMs or functional properties of normally-expressed enzymes may be impaired. The majority of enzymes examined in this report are themselves glycosylated; hence, the altered expression of these enzymes may not only affect the function of a variety of substrate proteins and lipids, but also the efficacy and kinetics of the enzyme-mediated glycosylation pathways in which they participate. Additional studies that examine enzyme protein expression in specific subcellular compartments, in neuronal versus glial cellular subpopulations, and/or in multiple brain regions in schizophrenia represent a possible avenue of future investigation. Studies of enzyme activity, kinetics, and upstream regulatory pathways may also provide insight into the relationship between dysregulated glycosylation enzyme transcription and abnormal neuroglycobiology in schizophrenia.

The current study illustrates that transcript expression of enzymes from multiple glycosylation pathways is considerably altered in schizophrenia, and that these alterations are not exclusive to ER-localized enzymes. Enzymes associated with both the addition and removal of glycans demonstrate increased mRNA expression in this illness and add to the growing body of evidence that protein PTMs are dysfunctional in schizophrenia brain. Wide-spread glycosylation abnormalities as a result of abnormal glycosylation enzyme expression could explain sex-specific neurodevelopmental alterations, deficits in neuritogenesis and abnormal dendritic spine morphology, impaired synaptic plasticity, altered protein localization, and many other substrate-specific deficits in schizophrenia. As such, dysglycosylation of substrates could represent a common impaired mechanism in multiple neurotransmitter systems and cell types. Future examinations of the neuroglycobiology of schizophrenia will be necessary to elucidate specific glycosides and glycan structures that contribute to the pathophysiology of this disorder.

## Acknowledgments

This work is supported by National Institutes of Health Grant MH53327 (JMW), MH064673 (VH), and MH066392 (VH).

## References

Acarin, L., Vela, J.M., González, B., Castellano, B., 1994. Demonstration of poly-N-acetyl lactosamine residues in ameboid and ramified microglial cells in rat brain by tomato lectin binding. J. Histochem. Cytochem. 42, 1033–41.

Albohy, A., Li, M.D., Zheng, R.B., Zou, C., Cairo, C.W., 2010. Insight into substrate recognition and catalysis by the human neuraminidase 3 (NEU3) through molecular modeling and site-directed mutagenesis. Glycobiology 20, 1127–1138. doi:10.1093/glycob/cwq077

Alghamdi, F., Guo, M., Abdulkhalek, S., Crawford, N., Amith, S.R., Szewczuk, M.R., 2014. A novel insulin receptor-signaling platform and its link to insulin resistance and type 2 diabetes. Cell. Signal. 26, 1355–1368. doi:10.1016/j.cellsig.2014.02.015

Baboval, T., Smith, F.I., 2002. Comparison of human and mouse Fuc-TX and Fuc-TXI genes, and expression studies in the mouse. Mamm. Genome 13, 538–41. doi:10.1007/s00335-001-2152-5

Barbeau, D., Liang, J.J., Robitalille, Y., Quirion, R., Srivastava, L.K., 1995. Decreased expression of the embryonic form of the neural cell adhesion molecule in schizophrenic brains. Proc. Natl. Acad. Sci. U. S. A. 92, 2785–9.

Bauer, D.E., Haroutunian, V., Meador-Woodruff, J.H., McCullumsmith, R.E., 2010. Abnormal glycosylation of EAAT1 and EAAT2 in prefrontal cortex of elderly patients with schizophrenia. Schizophr. Res. 117, 92–8. doi:10.1016/j.schres.2009.07.025

Bayer, T.A., Falkai, P., Maier, W., 1999. Genetic and non-genetic vulnerability factors in schizophrenia: the basis of the “two hit hypothesis”. J. Psychiatr. Res. 33, 543–8. doi:10.1016/S0022-3956(99)00039-4

Benjamini, Y., Hochberg, Y., 1995. Controlling the false discovery rate: a practical and powerful approach to multiple testing. J. R. Stat. Soc. 57, 289–300.

Bennett, E.P., Hassan, H., Hollingsworth, M.A., Clausen, H., 1999. A novel human UDP-N-acetyl-D-galactosamine:polypeptide N-acetylgalactosaminyltransferase, GalNAc-T7, with specificity for partial GalNAc-glycosylated acceptor substrates. FEBS Lett. 460, 226–30.

Bennett, E.P., Hassan, H., Mandel, U., Mirgorodskaya, E., Roepstorff, P., Burchell, J., Taylor-Papadimitriou, J., Hollingsworth, M.A., Merkx, G., van Kessel, A.G., Eiberg, H., Steffensen, R., Clausen, H., 1998. Cloning of a human UDP-N-acetyl-alpha-D-Galactosamine:polypeptide N-acetylgalactosaminyltransferase that complements other GalNAc-transferases in complete O-glycosylation of the MUC1 tandem repeat. J. Biol. Chem. 273, 30472–81.

Berezovska, O., McLean, P., Knowles, R., Frosh, M., Lu, F.M., Lux, S.E., Hyman, B.T., 1999. Notch1 inhibits neurite outgrowth in postmitotic primary neurons. Neuroscience 93, 433–439. doi:10.1016/S0306- 4522(99)00157-8

Berretta, S., 2012. Extracellular matrix abnormalities in schizophrenia. Neuropharmacology 62, 1584–1597. doi:10.1016/j.neuropharm.2011.08.010

Binder, J.X., Pletscher-Frankild, S., Tsafou, K., Stolte, C., O’Donoghue, S.I., Schneider, R., Jensen, L.J., 2014. COMPARTMENTS: unification and visualization of protein subcellular localization evidence. Database (Oxford). 2014, bau012. doi:10.1093/database/bau012

Bonten, E., van der Spoel, A., Fornerod, M., Grosveld, G., D’Azzo, A., 1996. Characterization of human lysosomal neuraminidase defines the molecular basis of the metabolic storage disorder sialidosis. Genes Dev. 10, 3156–69.

Butkinaree, C., Park, K., Hart, G.W., 2010. O-linked beta-N-acetylglucosamine (O-GlcNAc): Extensive crosstalk with phosphorylation to regulate signaling and transcription in response to nutrients and stress. Biochim. Biophys. Acta 1800, 96–106. doi:10.1016/j.bbagen.2009.07.018

Coyle, J.T., 1996. The glutamatergic dysfunction hypothesis for schizophrenia. Harv. Rev. Psychiatry 3, 241– 53.

d’Azzo, A., Bonten, E., 2010. Molecular mechanisms of pathogenesis in a glycosphingolipid and a glycoprotein storage disease. Biochem. Soc. Trans. 38, 1453–7. doi:10.1042/BST0381453

Davis, K.L., Kahn, R.S., Ko, G., Davidson, M., 1991. Dopamine in schizophrenia: A review and reconceptualization. Am. J. Psychiatry 148, 1474–1486. doi:1681750

DeMarco, M.L., Woods, R.J., 2008. Structural glycobiology: A game of snakes and ladders. Glycobiology. doi:10.1093/glycob/cwn026

Dennis, J.W., Lau, K.S., Demetriou, M., Nabi, I.R., 2009. Adaptive regulation at the cell surface by N-glycosylation. Traffic 10, 1569–78. doi:10.1111/j.1600-0854.2009.00981.x

Fritz, T.A., Raman, J., Tabak, L.A., 2006. Dynamic association between the catalytic and lectin domains of human UDP-GalNAc:polypeptide alpha-N-acetylgalactosaminyltransferase-2. J. Biol. Chem. 281, 8613–9. doi:10.1074/jbc.M513590200

Fukuda, T., Hashimoto, H., Okayasu, N., Kameyama, A., Onogi, H., Nakagawasai, O., Nakazawa, T., Kurosawa, T., Hao, Y., Isaji, T., Tadano, T., Narimatsu, H., Taniguchi, N., Gu, J., 2011. alpha-1,6-fucosyltransferase-deficient mice exhibit multiple behavioral abnormalities associated with a schizophrenia-like phenotype: Importance of the balance between the dopamine and serotonin systems. J. Biol. Chem. 286, 18434–18443. doi:10.1074/jbc.M110.172536

Giordanengo, V., Bannwarth, S., Laffont, C., Miecem, V., Harduin-Lepers, A., Delannoy, P., Lefebvre, J.-C., 1997. Cloning and Expression of cDNA for a Human Gal(beta1-3)GalNAc alpha2,3-Sialyltransferase from the CEM T-Cell Line. Eur. J. Biochem. 247, 558–566. doi:10.1111/j.1432-1033.1997.00558.x

Gottschall, P.E., Howell, M.D., 2015. ADAMTS expression and function in central nervous system injury and disorders. Matrix Biol. 44–46, 70–6. doi:10.1016/j.matbio.2015.01.014

Gu, W., Fukuda, T., Isaji, T., Hang, Q., Lee, H., Sakai, S., Morise, J., Mitoma, J., Higashi, H., Taniguchi, N., Yawo, H., Oka, S., Gu, J., 2015. Loss of α1,6-Fucosyltransferase Decreases Hippocampal Long Term Potentiation: IMPLICATIONS FOR CORE FUCOSYLATION IN THE REGULATION OF AMPA RECEPTOR HETEROMERIZATION AND CELLULAR SIGNALING. J. Biol. Chem. 290, 17566–17575. doi:10.1074/jbc.M114.579938

Gu, W., Fukuda, T., Isaji, T., Hashimoto, H., Wang, Y., Gu, J., 2013. α1,6-Fucosylation regulates neurite formation via the activin/phospho-Smad2 pathway in PC12 cells: the implicated dual effects of Fut8 for TGF-β/activin-mediated signaling. FASEB J. 27, 3947–58. doi:10.1096/fj.12-225805

Guo, S., Sato, T., Shirane, K., Furukawa, K., 2001. Galactosylation of N-linked oligosaccharides by human beta-1,4-galactosyltransferases I, II, III, IV, V, and VI expressed in Sf-9 cells. Glycobiology 11, 813–820.

Häfner, H., Maurer, K., Löffler, W., Fätkenheuer, B., an der Heiden, W., Riecher-Rössler, A., Behrens, S., Gattaz, W.F., 1994. The epidemiology of early schizophrenia. Influence of age and gender on onset and early course. Br. J. Psychiatry. Suppl. 164, 29–38.

Hammond, J.C., McCullumsmith, R.E., Funk, A.J., Haroutunian, V., Meador-Woodruff, J.H., 2010. Evidence for abnormal forward trafficking of AMPA receptors in frontal cortex of elderly patients with schizophrenia. Neuropsychopharmacology 35, 2110–9. doi:10.1038/npp.2010.87

Hammond, J.C., McCullumsmith, R.E., Haroutunian, V., Meador-Woodruff, J.H., 2011. Endosomal trafficking of AMPA receptors in frontal cortex of elderly patients with schizophrenia. Schizophr. Res. 130, 260–5. doi:10.1016/j.schres.2011.04.029

Harris, R.J., Spellman, M.W., 1993. O-linked fucose and other post-translational modifications unique to EGF modules. Glycobiology 3, 219–24.

Harrison, P.J., 2011. Using our brains: the findings, flaws, and future of postmortem studies of psychiatric disorders. Biol. Psychiatry 69, 102–3. doi:10.1016/j.biopsych.2010.09.008

Harrison, P.J., 2000. Postmortem studies in schizophrenia. Dialogues Clin. Neurosci. 2, 349–57.

Harrison, P.J., Weinberger, D.R., 2005. Schizophrenia genes, gene expression, and neuropathology: on the matter of their convergence. Mol. Psychiatry 10, 40–68. doi:10.1038/sj.mp.4001558

Heal, D.J., Gosden, J., Jackson, H.C., Cheetham, S.C., Smith, S.L., 2012. Metabolic Consequences of Antipsychotic Therapy: Preclinical and Clinical Perspectives on Diabetes, Diabetic Ketoacidosis, and Obesity, in: Gross, G., Geyer, M.A. (Eds.), Current Antipsychotics, Handbook of Experimental Pharmacology. Springer Berlin Heidelberg, Berlin, Heidelberg, pp. 135–164. doi:10.1007/978-3-642-25761-2

Helenius, A., Aebi, M., 2004. Roles of N-linked glycans in the endoplasmic reticulum. Annu. Rev. Biochem. 73, 1019–49. doi:10.1146/annurev.biochem.73.011303.073752

Howerton, C.L., Morgan, C.P., Fischer, D.B., Bale, T.L., 2013. O-GlcNAc transferase (OGT) as a placental biomarker of maternal stress and reprogramming of CNS gene transcription in development. Proc. Natl. Acad. Sci. U. S. A. 110, 5169–74. doi:10.1073/pnas.1300065110

Ikemoto, K., 2014. Lectin-Positive Spherical Deposits (SPD) Detected in the Molecular Layer of Hippocampal Dentate Gyrus of Dementia, Down’s Syndrome, and Schizophrenia. J. Alzheimer’s Dis. Park. 04, 4–7. doi:10.4172/2161-0460.1000169

Irvine, K.D., 2008. A notch sweeter. Cell 132, 177–9. doi:10.1016/j.cell.2008.01.005

Ishida, H., Togayachi, A., Sakai, T., Iwai, T., Hiruma, T., Sato, T., Okubo, R., Inaba, N., Kudo, T., Gotoh, M., Shoda, J., Tanaka, N., Narimatsu, H., 2005. A novel beta1,3-N-acetylglucosaminyltransferase (beta3Gn-T8), which synthesizes poly-N-acetyllactosamine, is dramatically upregulated in colon cancer. FEBS Lett. 579, 71–8. doi:10.1016/j.febslet.2004.11.037

Isomura, R., Kitajima, K., Sato, C., 2011. Structural and functional impairments of polysialic acid by a mutated polysialyltransferase found in schizophrenia. J. Biol. Chem. 286, 21535–45. doi:10.1074/jbc.M111.221143

Kato, K., Shiga, K., Yamaguchi, K., Hata, K., Kobayashi, T., Miyazaki, K., Saijo, S., Miyagi, T., 2006. Plasma-membrane-associated sialidase (NEU3) differentially regulates integrin-mediated cell proliferation through laminin- and fibronectin-derived signalling. Biochem. J. 394, 647–656. doi:10.1042/BJ20050737

Keith, N., Parodi, A.J., Caramelo, J.J., 2005. Glycoprotein tertiary and quaternary structures are monitored by the same quality control mechanism. J. Biol. Chem. 280, 18138–18141. doi:10.1074/jbc.M501710200

Khandaker, G.M., Zimbron, J., Lewis, G., Jones, P.B., 2013. Prenatal maternal infection, neurodevelopment and adult schizophrenia : a systematic review of population-based studies. Psychol. Med. 43, 239–57. doi:10.1017/S0033291712000736

Kim, D.H., Stahl, S.M., 2010. Behavioral Neurobiology of Schizophrenia and Its Treatment, in: Swerdlow, N.R. (Ed.), Behavioral Neurobiology of Schizophrenia and Its Treatment, Current Topics in Behavioral Neurosciences. Springer Berlin Heidelberg, Berlin, Heidelberg, pp. 123–139. doi:10.1007/978-3-642-13717-4

Kim, P., Scott, M.R., Meador-Woodruff, J.H., 2018. Abnormal expression of ER quality control and ER associated degradation proteins in the dorsolateral prefrontal cortex in schizophrenia. Schizophr. Res. doi:10.1016/j.schres.2018.02.010

Kippe, J.M., Mueller, T.M., Haroutunian, V., Meador-Woodruff, J.H., 2015. Abnormal N-acetylglucosaminyltransferase expression in prefrontal cortex in schizophrenia. Schizophr. Res. 166, 219–24. doi:10.1016/j.schres.2015.06.002

Kötzler, M.P., Blank, S., Bantleon, F.I., Spillner, E., Meyer, B., 2012. Donor substrate binding and enzymatic mechanism of human core α1,6-fucosyltransferase (FUT8). Biochim. Biophys. Acta 1820, 1915–25. doi:10.1016/j.bbagen.2012.08.018

Kristiansen, L. V, Huerta, I., Beneyto, M., Meador-Woodruff, J.H., 2007. NMDA receptors and schizophrenia. Curr. Opin. Pharmacol. 7, 48–55. doi:10.1016/j.coph.2006.08.013

Kuan, C.-T., Chang, J., Mansson, J.-E., Li, J., Pegram, C., Fredman, P., McLendon, R.E., Bigner, D.D., 2010. Multiple phenotypic changes in mice after knockout of the B3gnt5 gene, encoding Lc3 synthase--a key enzyme in lacto-neolacto ganglioside synthesis. BMC Dev. Biol. 10, 114. doi:10.1186/1471-213X-10-114

Kumar, R., Yang, J., Larsen, R.D., Stanley, P., 1990. Cloning and expression of N-acetylglucosaminyltransferase I, the medial Golgi transferase that initiates complex N-linked carbohydrate formation. Proc. Natl. Acad. Sci. U. S. A. 87, 9948–52.

Kurimoto, A., Kitazume, S., Kizuka, Y., Nakajima, K., Oka, R., Fujinawa, R., Korekane, H., Yamaguchi, Y., Wada, Y., Taniguchi, N., 2014. The Absence of Core Fucose Up-regulates GnT-III and Wnt Target Genes: A POSSIBLE MECHANISM FOR AN ADAPTIVE RESPONSE IN TERMS OF GLYCAN FUNCTION. J. Biol. Chem. 289, 11704–11714. doi:10.1074/jbc.M113.502542

Lai, E.C., 2004. Notch signaling: control of cell communication and cell fate. Development 131, 965–973. doi:10.1242/dev.01074

Livak, K.J., Schmittgen, T.D., 2001. Analysis of relative gene expression data using real-time quantitative PCR and the 2(-Delta Delta C(T)) Method. Methods 25, 402–8. doi:10.1006/meth.2001.1262

Maccioni, H.J.F., Quiroga, R., Ferrari, M.L., 2011. Cellular and molecular biology of glycosphingolipid glycosylation. J. Neurochem. 117, 589–602. doi:10.1111/j.1471-4159.2011.07232.x

Magesh, S., Suzuki, T., Miyagi, T., Ishida, H., Kiso, M., 2006. Homology modeling of human sialidase enzymes NEU1, NEU3 and NEU4 based on the crystal structure of NEU2: Hints for the design of selective NEU3 inhibitors. J. Mol. Graph. Model. 25, 196–207. doi:10.1016/j.jmgm.2005.12.006

McCullumsmith, R.E., Clinton, S.M., Meador-Woodruff, J.H., 2003. Schizophrenia as a disorder of neuroplasticity. Int. Rev. Neurobiol. doi:10.1016/S0074-7742(04)59002-5

McCullumsmith, R.E., Meador-Woodruff, J.H., 2011. Novel approaches to the study of postmortem brain in psychiatric illness: old limitations and new challenges. Biol. Psychiatry 69, 127–33. doi:10.1016/j.biopsych.2010.09.035

McGuire, J.L., Depasquale, E.A., Funk, A.J., O’Donnovan, S.M., Hasselfeld, K., Marwaha, S., Hammond, J.H., Hartounian, V., Meador-Woodruff, J.H., Meller, J., McCullumsmith, R.E., 2017. Abnormalities of signal transduction networks in chronic schizophrenia. npj Schizophr. 3, 30. doi:10.1038/s41537-017-0032-6

McGuire, J.L., Hammond, J.H., Yates, S.D., Chen, D., Haroutunian, V., Meador-Woodruff, J.H., McCullumsmith, R.E., 2014. Altered serine/threonine kinase activity in schizophrenia. Brain Res. 1568, 42–54. doi:10.1016/j.brainres.2014.04.029

Meador-Woodruff, J.H., Healy, D.J., 2000. Glutamate receptor expression in schizophrenic brain, in: Brain Research Reviews. pp. 288–294. doi:10.1016/S0165-0173(99)00044-2

Miyagi, T., Wada, T., Yamaguchi, K., Shiozaki, K., Sato, I., Kakugawa, Y., Yamanami, H., Fujiya, T., 2008. Human sialidase as a cancer marker. Proteomics 8, 3303–3311. doi:10.1002/pmic.200800248

Mollicone, R., Moore, S.E.H., Bovin, N., Garcia-Rosasco, M., Candelier, J.-J., Martinez-Duncker, I., Oriol, R., 2009. Activity, splice variants, conserved peptide motifs, and phylogeny of two new alpha1,3- fucosyltransferase families (FUT10 and FUT11). J. Biol. Chem. 284, 4723–38. doi:10.1074/jbc.M809312200

Moon, S.K., Cho, S.H., Kim, K.W., Jeon, J.H., Ko, J.H., Kim, B.Y., Kim, C.H., 2007. Overexpression of membrane sialic acid-specific sialidase Neu3 inhibits matrix metalloproteinase-9 expression in vascular smooth muscle cells. Biochem. Biophys. Res. Commun. 356, 542–547. doi:10.1016/j.bbrc.2007.02.155

Moremen, K.W., Nairn, A. V, 2014. Handbook of Glycosyltransferases and Related Genes, 2nd ed, Handbook of Glycosyltransferases and Related Genes. Springer Japan, Tokyo. doi:10.1007/978-4-431-54240-7

Moremen, K.W., Tiemeyer, M., Nairn, A. V, 2012. Vertebrate protein glycosylation: diversity, synthesis and function. Nat. Rev. Mol. Cell Biol. 13, 448–62. doi:10.1038/nrm3383

Motulsky, H.J., Brown, R.E., 2006. Detecting outliers when fitting data with nonlinear regression - a new method based on robust nonlinear regression and the false discovery rate. BMC Bioinformatics 7, 123. doi:10.1186/1471-2105-7-123

Mueller, T.M., Haroutunian, V., Meador-Woodruff, J.H., 2014. N-glycosylation of GABAA receptor subunits is altered in schizophrenia. Neuropsychopharmacology 39, 528–37. doi:10.1038/npp.2013.190

Mueller, T.M., Remedies, C.E., Haroutunian, V., Meador-Woodruff, J.H., 2015. Abnormal subcellular localization of GABAA receptor subunits in schizophrenia brain. Transl. Psychiatry 5, e612. doi:10.1038/tp.2015.102

Mueller, T.M., Yates, S.D., Haroutunian, V., Meador-Woodruff, J.H., 2017. Altered fucosyltransferase expression in the superior temporal gyrus of elderly patients with schizophrenia. Schizophr. Res. 182, 66–73. doi:10.1016/j.schres.2016.10.024

Nakazawa, K., Zsiros, V., Jiang, Z., Nakao, K., Kolata, S., Zhang, S., Belforte, J.E., 2012. GABAergic interneuron origin of schizophrenia pathophysiology. Neuropharmacology 62, 1574–83. doi:10.1016/j.neuropharm.2011.01.022

Narayan, S., Head, S.R., Gilmartin, T.J., Dean, B., Thomas, E.A., 2009. Evidence for disruption of sphingolipid metabolism in schizophrenia. J. Neurosci. Res. 87, 278–288. doi:10.1002/jnr.21822

Niimi, K., Nishioka, C., Miyamoto, T., Takahashi, E., Miyoshi, I., Itakura, C., Yamashita, T., 2011. Impairment of neuropsychological behaviors in ganglioside GM3-knockout mice. Biochem. Biophys. Res. Commun. 406, 524–8. doi:10.1016/j.bbrc.2011.02.071

Ohtsubo, K., Marth, J.D., 2006. Glycosylation in cellular mechanisms of health and disease. Cell 126, 855–67. doi:10.1016/j.cell.2006.08.019

Olivari, S., Molinari, M., 2007. Glycoprotein folding and the role of EDEM1, EDEM2 and EDEM3 in degradation of folding-defective glycoproteins. FEBS Lett. 581, 3658–64. doi:10.1016/j.febslet.2007.04.070

Pantazopoulos, H., Boyer-Boiteau, A., Holbrook, E.H., Jang, W., Hahn, C.G., Arnold, S.E., Berretta, S., 2013. Proteoglycan abnormalities in olfactory epithelium tissue from subjects diagnosed with schizophrenia. Schizophr. Res. 150, 366–372. doi:10.1016/j.schres.2013.08.013

Pietersen, C.Y., Mauney, S. a, Kim, S.S., Passeri, E., Lim, M.P., Rooney, R.J., Goldstein, J.M., Petreyshen, T.L., Seidman, L.J., Shenton, M.E., Mccarley, R.W., Sonntag, K.-C., Woo, T.-U.W., Petryshen, T.L., Seidman, L.J., Shenton, M.E., Mccarley, R.W., Sonntag, K.-C., Woo, T.-U.W., 2014. Molecular profiles of parvalbumin-immunoreactive neurons in the superior temporal cortex in schizophrenia. J. Neurogenet. 28, 70–85. doi:10.3109/01677063.2013.878339

Powchik, P., Davidson, M., Nemeroff, C.B., Haroutunian, V., Purohit, D., Losonczy, M., Bissette, G., Perl, D., Ghanbari, H., Miller, B., 1993. Alzheimer’s-disease-related protein in geriatric schizophrenic patients with cognitive impairment. Am. J. Psychiatry 150, 1726–7. doi:10.1176/ajp.150.11.1726

Purohit, D.P., Perl, D.P., Haroutunian, V., Powchik, P., Davidson, M., Davis, K.L., 1998. Alzheimer disease and related neurodegenerative diseases in elderly patients with schizophrenia: a postmortem neuropathologic study of 100 cases. Arch. Gen. Psychiatry 55, 205–211.

Redmond, L., Oh, S.R., Hicks, C., Weinmaster, G., Ghosh, A., 2000. Nuclear Notch1 signaling and the regulation of dendritic development. Nat. Neurosci. 3, 30–40. doi:10.1038/71104

Ricketts, L.M., Dlugosz, M., Luther, K.B., Haltiwanger, R.S., Majerus, E.M., 2007. O-fucosylation is required for ADAMTS13 secretion. J. Biol. Chem. 282, 17014–23. doi:10.1074/jbc.M700317200

Roth, J., Zuber, C., Park, S., Jang, I., Lee, Y., Kysela, K.G., Le Fourn, V., Santimaria, R., Guhl, B., Cho, J.W., 2010. Protein N-glycosylation, protein folding, and protein quality control. Mol. Cells 30, 497–506. doi:10.1007/s10059-010-0159-z

Röttger, S., White, J., Wandall, H.H., Olivo, J.C., Stark, A., Bennett, E.P., Whitehouse, C., Berger, E.G., Clausen, H., Nilsson, T., 1998. Localization of three human polypeptide GalNAc-transferases in HeLa cells suggests initiation of O-linked glycosylation throughout the Golgi apparatus. J. Cell Sci. 111, 45–60.

Rubio, M.D., Wood, K., Haroutunian, V., Meador-Woodruff, J.H., 2013. Dysfunction of the ubiquitin proteasome and ubiquitin-like systems in schizophrenia. Neuropsychopharmacology 38, 1910–20. doi:10.1038/npp.2013.84

Ryšlavá, H., Doubnerová, V., Kavan, D., Vaněk, O., 2013. Effect of posttranslational modifications on enzyme function and assembly. J. Proteomics. doi:10.1016/j.jprot.2013.03.025

Sato, C., Kitajima, K., 2013. Impact of structural aberrancy of polysialic acid and its synthetic enzyme ST8SIA2 in schizophrenia. Front. Cell. Neurosci. 7, 61. doi:10.3389/fncel.2013.00061

Sato, T., Sato, M., Kiyohara, K., Sogabe, M., Shikanai, T., Kikuchi, N., Togayachi, A., Ishida, H., Ito, H., Kameyama, A., Gotoh, M., Narimatsu, H., 2006. Molecular cloning and characterization of a novel human beta1,3-glucosyltransferase, which is localized at the endoplasmic reticulum and glucosylates O-linked fucosylglycan on thrombospondin type 1 repeat domain. Glycobiology 16, 1194–1206. doi:cwl035[pii]\r10.1093/glycob/cwl035

Scaringi, R., Piccoli, M., Papini, N., Cirillo, F., Conforti, E., Bergante, S., Tringali, C., Garatti, A., Gelfi, C., Venerando, B., Menicanti, L., Tettamanti, G., Anastasia, L., 2013. NEU3 sialidase is activated under hypoxia and protects skeletal muscle cells from apoptosis through the activation of the epidermal growth factor receptor signaling pathway and the hypoxia-inducible factor (HIF)-1α. J. Biol. Chem. 288, 3153–62. doi:10.1074/jbc.M112.404327

Schachter, H., 1986. Biosynthetic controls that determine the branching and microheterogeneity of protein-bound oligosaccharides. Biochem. Cell Biol. 64, 163–81.

Schnaar, R.L., Gerardy-Schahn, R., Hildebrandt, H., 2014. Sialic acids in the brain: gangliosides and polysialic acid in nervous system development, stability, disease, and regeneration. Physiol. Rev. 94, 461–518. doi:10.1152/physrev.00033.2013

Scott, M.R., Rubio, M.D., Haroutunian, V., Meador-Woodruff, J.H., 2015. Protein Expression of Proteasome Subunits in Elderly Patients with Schizophrenia. Neuropsychopharmacology. doi:10.1038/npp.2015.219

Seko, A., Yamashita, K., 2005. Characterization of a novel galactose beta1,3-N-acetylglucosaminyltransferase (beta3Gn-T8): the complex formation of beta3Gn-T2 and beta3Gn-T8 enhances enzymatic activity. Glycobiology 15, 943–51. doi:10.1093/glycob/cwi082

Shayevitz, C., Cohen, O.S., Faraone, S. V, Glatt, S.J., 2012. A re-review of the association between the NOTCH4 locus and schizophrenia. Am. J. Med. Genet. Part B Neuropsychiatr. Genet. 159B, 477–483. doi:10.1002/ajmg.b.32050

Shiraishi, N., Natsume, A., Togayachi, A., Endo, T., Akashima, T., Yamada, Y., Imai, N., Nakagawa, S., Koizumi, S., Sekine, S., Narimatsu, H., Sasaki, K., 2001. Identification and characterization of three novel beta 1,3-N-acetylglucosaminyltransferases structurally related to the beta 1,3-galactosyltransferase family. J. Biol. Chem. 276, 3498–507. doi:10.1074/jbc.M004800200

Sobkowicz, A.D., Gallagher, M.E., Reid, C.J., Crean, D., Carrington, S.D., Irwin, J.A., 2014. Modulation of expression in BEAS-2B airway epithelial cells of α-L-fucosidase A1 and A2 by Th1 and Th2 cytokines, and overexpression of α-L-fucosidase 2. Mol. Cell. Biochem. 390, 101–13. doi:10.1007/s11010-014-1961-2

Spiro, R.G., 2002. Protein glycosylation: nature, distribution, enzymatic formation, and disease implications of glycopeptide bonds. Glycobiology 12, 43R–56R. doi:10.1093/glycob/12.4.43R

Stanta, J.L., Saldova, R., Struwe, W.B., Byrne, J.C., Leweke, F.M., Rothermund, M., Rahmoune, H., Levin, Y., Guest, P.C., Bahn, S., Rudd, P.M., 2010. Identification of N-glycosylation changes in the CSF and serum in patients with schizophrenia. J. Proteome Res. 9, 4476–89. doi:10.1021/pr1002356

Takasaki, N., Tachibana, K., Ogasawara, S., Matsuzaki, H., Hagiuda, J., Ishikawa, H., Mochida, K., Inoue, K., Ogonuki, N., Ogura, A., Noce, T., Ito, C., Toshimori, K., Narimatsu, H., 2014. A heterozygous mutation of GALNTL5 affects male infertility with impairment of sperm motility. Proc. Natl. Acad. Sci. U. S. A. 111, 1120–5. doi:10.1073/pnas.1310777111

Telford, J.E., Bones, J., McManus, C., Saldova, R., Manning, G., Doherty, M., Leweke, F.M., Rothermundt, M., Guest, P.C., Rahmoune, H., Bahn, S., Rudd, P.M., 2012. Antipsychotic treatment of acute paranoid schizophrenia patients with olanzapine results in altered glycosylation of serum glycoproteins. J. Proteome Res. 11, 3743–52. doi:10.1021/pr300218h

The UniProt Consortium, 2014. UniProt: a hub for protein information. Nucleic Acids Res. 43, D204–D212. doi:10.1093/nar/gku989

Tringali, C., Lupo, B., Cirillo, F., Papini, N., Anastasia, L., Lamorte, G., Colombi, P., Bresciani, R., Monti, E., Tettamanti, G., Venerando, B., 2009. Silencing of membrane-associated sialidase Neu3 diminishes apoptosis resistance and triggers megakaryocytic differentiation of chronic myeloid leukemic cells K562 through the increase of ganglioside GM3. Cell Death Differ. 16, 164–174. doi:10.1038/cdd.2008.141

Tucholski, J., Simmons, M.S., Pinner, A.L., Haroutunian, V., McCullumsmith, R.E., Meador-Woodruff, J.H., 2013a. Abnormal N-linked glycosylation of cortical AMPA receptor subunits in schizophrenia. Schizophr. Res. 146, 177–83. doi:10.1016/j.schres.2013.01.031

Tucholski, J., Simmons, M.S., Pinner, A.L., McMillan, L.D., Haroutunian, V., Meador-Woodruff, J.H., 2013b. N-linked glycosylation of cortical N-methyl-D-aspartate and kainate receptor subunits in schizophrenia. Neuroreport 24, 688–91. doi:10.1097/WNR.0b013e328363bd8a

Vanhooren, V., Dewaele, S., Kuro-O, M., Taniguchi, N., Dollé, L., van Grunsven, L.A., Makrantonaki, E., Zouboulis, C.C., Chen, C.C., Libert, C., 2011. Alteration in N-glycomics during mouse aging: a role for FUT8. Aging Cell 10, 1056–66. doi:10.1111/j.1474-9726.2011.00749.x

Varki, A., Cummings, R.D., Esko, J.D., Freeze, H.H., Stanley, P., Bertozzi, C.R., Hart, G.W., Etzler, M.E., 2009. Essentials of Glycobiology, 2nd ed, Essentials of Glycobiology. Cold Spring Harbor Laboratory Press.

Wang, L.W., Dlugosz, M., Somerville, R.P.T., Raed, M., Haltiwanger, R.S., Apte, S.S., 2007. O-fucosylation of thrombospondin type 1 repeats in ADAMTS-like-1/punctin-1 regulates secretion: implications for the ADAMTS superfamily. J. Biol. Chem. 282, 17024–31. doi:10.1074/jbc.M701065200

Wang, X., Gu, J., Ihara, H., Miyoshi, E., Honke, K., Taniguchi, N., 2006. Core fucosylation regulates epidermal growth factor receptor-mediated intracellular signaling. J. Biol. Chem. 281, 2572–7. doi:10.1074/jbc.M510893200

Wang, X., Inoue, S., Gu, J., Miyoshi, E., Noda, K., Li, W., Mizuno-Horikawa, Y., Nakano, M., Asahi, M., Takahashi, M., Uozumi, N., Ihara, S., Lee, S.H., Ikeda, Y., Yamaguchi, Y., Aze, Y., Tomiyama, Y., Fujii, J., Suzuki, K., Kondo, A., Shapiro, S.D., Lopez-Otin, C., Kuwaki, T., Okabe, M., Honke, K., Taniguchi, N., 2005. Dysregulation of TGF-beta1 receptor activation leads to abnormal lung development and emphysema-like phenotype in core fucose-deficient mice. Proc. Natl. Acad. Sci. U. S. A. 102, 15791–6. doi:10.1073/pnas.0507375102

Wang, Y., Shao, L., Shi, S., Harris, R.J., Spellman, M.W., Stanley, P., Haltiwanger, R.S., 2001. Modification of epidermal growth factor-like repeats with O-fucose. Molecular cloning and expression of a novel GDP-fucose protein O-fucosyltransferase. J. Biol. Chem. 276, 40338–45. doi:10.1074/jbc.M107849200

Yang, X.-X., Zhu, A.-N., Li, F.-X., Zhang, Z.-X., Li, M., 2013. Neurogenic locus notch homolog protein 4 and brain-derived neurotrophic factor variants combined effect on schizophrenia susceptibility. Acta Neuropsychiatr. 25, 356–60. doi:10.1017/neu.2013.13

Yip, B., Chen, S.H., Mulder, H., Höppener, J.W., Schachter, H., 1997. Organization of the human beta-1,2-N-acetylglucosaminyltransferase I gene (MGAT1), which controls complex and hybrid N-glycan synthesis. Biochem. J. 321 ( Pt 2, 465–74.

Yogalingam, G., Bonten, E.J., van de Vlekkert, D., Hu, H., Moshiach, S., Connell, S.A., D’Azzo, A., 2008. Neuraminidase 1 is a negative regulator of lysosomal exocytosis. Dev. Cell 15, 74–86. doi:10.1016/j.devcel.2008.05.005

Yu, R.K., Tsai, Y.-T., Ariga, T., Yanagisawa, M., 2011. Structures, biosynthesis, and functions of gangliosides--an overview. J. Oleo Sci. 60, 537–44.

